# ALBIS: simulation of multi-resolution and multi-dimensional spatial transcriptomics data

**DOI:** 10.64898/2026.10.06.757026

**Authors:** Jan Matthias, Jianing Yao, Stephanie C. Hicks

**Author notes:** These authors contributed equally (Jan Matthias, Jianing Yao).

## Abstract

The rapid adoption of spatial transcriptomics (ST) technologies has been accompanied by substantial growth in computational methods, creating an increasing need for systematic and reliable benchmarking, particularly as analyses increasingly extend from individual tissue sections to three-dimensional (3D) tissue reconstruction. Existing benchmarks often rely on a small and similar set of experimentally-derived datasets because they offer a reliable ground truth label, which is hard to find in practice. Simulated or synthetic data provide an alternative, yet most existing simulators either adapt single-cell RNA sequencing models to spatial coordinates or generate individual sections in a two-dimensional (2D) space at predefined resolutions. More recent approaches can simulate 2D and 3D tissues, but generally do not derive multiple spatial resolutions from the same molecular realization. Here, we introduce ALBIS (A mu**L**ti-resolution **B**iological **I**n-silica **S**imulator for spatial transcriptomics), a statistically grounded and highly customizable framework that generates synthetic ST data from an *in silica* 3D tissue. ALBIS defines spatially structured cell populations with corresponding markers for gene expression, then initializes individual mRNA molecules in 3D space to create cell-level information with a ground truth 3D coordinate for each molecule and cell. From this common realization, ALBIS cuts the 3D tissue into multiple 2D tissue sections with cell-, bin-, and spot-level measurements, enabling direct comparison across spatial resolutions. Properties of the simulated data, including spatial domains, cell type composition, gene-expression effects, batch effects, measurement resolution, and tissue scale, can be systematically varied, enabling rigorous stress-testing of computational methods under known and increasingly challenging conditions.

## 1 Introduction

Spatial transcriptomics (ST) has enabled a new class of biological analyses by measuring gene expression while preserving the spatial context of the tissue [1]. The rapid adoption of these spatial profiling technologies has been accompanied by the development of a large and growing ecosystem of computational methods, with nearly 600 software tools available for downstream analyses as of 2026 [2]. As multiple methods are often proposed for the same data analysis task, systematic benchmarking has become essential to evaluate their relative performance and identify the conditions under which they succeed or fail. However, existing benchmarks frequently rely on a relatively small set of well-studied experimental datasets that are repeatedly used for method development and evaluation. Many of these datasets also lack reliable ground-truth labels, which complicates objective assessment of method performance and limits the ability to determine whether results generalize to less-studied datasets [3]. These challenges are even greater for serial-section alignment and 3D tissue reconstruction because experimental data generally do not provide the true spatial relationships among sections or their underlying 3D coordinates. Simulated data offer an alternative by providing known labels, spatial relationships, and coordinates under controlled biological and technical settings, and can support both method development and systematic benchmarking.

Current generative simulators for spatial transcriptomics differ in how spatial structure is incorporated into the data-generating process. Some approaches adapt existing single-cell RNA-sequencing (scRNA-seq) simulation frameworks to generate spatially resolved data, whereas native spatial simulators model spatial organization directly. Across both groups, simulators may also be *reference-based*, in which expression parameters, spatial patterns, or both are learned from a reference dataset, or *reference-free* [4,5], in which the spatial data-generating process is specified independently of a spatial reference.

Among approaches adapted from scRNA-seq simulation frameworks, widely used simulators such as Splatter [6], ZINB-WaVE [7], scDesign2 [8], SPARSim [9], and SymSim [10] provide established methods for generating synthetic gene-expression count data, but they do not explicitly model spatial coordinates or spatial relationships between observations. SimAdaptor [11] provides a strategy for adapting these non-spatial simulators to spatial transcriptomics data. It first partitions a reference spatial transcriptomics dataset into spatial regions using BayesSpace [12] and then independently applies existing expression simulators within each region, thereby incorporating the regional structure of the reference dataset into the simulated expression data.

Native spatial transcriptomics simulators instead incorporate spatial structure directly into the generative process. SRTsim [13] simulates gene-specific expression from a Poisson, zero-inflated Poisson (ZIP), negative binomial (NB), or zero-inflated negative binomial (ZINB) distribution and supports both reference-based and reference-free simulation, although reference-free spatial patterns must be specified by the user rather than learned from observed data. scDesign3 [14] uses generalized additive models with a copula-based framework to learn spatial expression patterns from reference data. However, neither framework explicitly constructs multiple tissue sections from a shared 3D spatial realization or models coordinated variation across serial slices. This limits their use for benchmarking methods that depend on known relationships across sections, including serial-section alignment, 3D tissue reconstruction, and cross-section integration [15–17].

More recent native spatial simulators, including scCube [17] and Spider [5], extend simulation to three-dimensional (3D) tissues. scCube can generate reference-free 3D spatial patterns and divide the resulting tissue into sequential 2D slices while allowing users to vary spatial resolution and spot arrangement. Spider similarly generates 2D or 3D cellular layouts from user-defined cell type proportions and spatial adjacency patterns and can subsequently aggregate single-cell expression profiles into spot-level observations. However, both frameworks represent expression at the cell level and derive coarser-resolution measurements by aggregating cell-expression profiles rather than explicitly simulating the spatial locations of individual transcripts. Thus, cross-resolution correspondence is defined through cell-level aggregation rather than capture from a shared spatial realization of individual RNA molecules.

Here, we introduce ALBIS **(A** mu**L**ti-resolution **B**iological **I**n-silico **S**imulator for spatial transcriptomics), a 3D simulation framework that generates multiple spatial measurement resolutions from a shared transcript-level molecular ground truth. ALBIS constructs a 3D tissue with domain-dependent cell type composition and individual RNA molecules with explicit spatial coordinates, and uses this common molecular realization to generate cellular-resolution (cell-level), subcellular-resolution (bin-level), and multicellular-resolution (spot-level) observations. Because all measurement scales are derived from the same underlying tissue, ALBIS provides direct ground-truth correspondence across resolutions together with known cell identities, spatial domains, and 3D coordinates. The simulated tissue can also be sectioned into serial slices and subjected to controlled technical variation, allowing ALBIS to support method development and benchmarking for spatial alignment, 3D reconstruction, cross-resolution integration, and other downstream spatial transcriptomics analyses.

In this study, we introduce our multi-resolution and multi-dimensional simulation framework and characterize the statistical properties of ALBIS-generated data through comparison with experimentally derived spatial transcriptomics datasets, including recovery of cell types and spatial domains as well as spatial deconvolution. We further demonstrate the utility of ALBIS across a range of benchmarking applications, including cross-resolution and cross-technology analyses and controlled stress testing of spatial methods.

## 2 Results

### 2.1 A multi-dimensional, multi-resolution simulator for spatial transcriptomics data

ALBIS is a multi-dimensional and multi-resolution framework for simulating spatial transcriptomics data at transcript and cellular scales, which consists of six sequential steps **(Figure 1A)**. First, we generate a 3D spherical tissue volume in which cells are placed. Second, cells are assigned to spatial domains. In the example shown here, the tissue contains five wedge-shaped domains surrounding a central core domain, although the number and geometry of domains can be modified by the user. Third, cell types are assigned conditional on spatial domain, allowing particular cell types to be enriched within specific regions while maintaining heterogeneous cell type composition. Then, gene expression counts are simulated for each cell using a negative binomial (NB) model, which has been shown to appropriately model the data-generating mechanism for spatial transcriptomics data [18−20]. These counts are then expanded into individual mRNA molecules, each with an associated gene identity and spatial coordinate. Finally, the simulated 3D coordinates at the molecule and cell level are converted into resolution-specific spatial transcriptomics datasets. The same underlying transcript-level ground truth is used to generate observations at multiple spatial resolutions: molecules can be aggregated into cells, rectangular bins, or circular spots to produce cellular-resolution, subcellular-resolution, and multicellular-resolution data, respectively. In the example shown **(Figure 1B)**, transcripts are aggregated into subcellular-resolution bins. The resulting sections are then independently rotated and translated to generate misaligned inputs while preserving the original spatial coordinates as ground truth for benchmarking.

**Figure 1:**
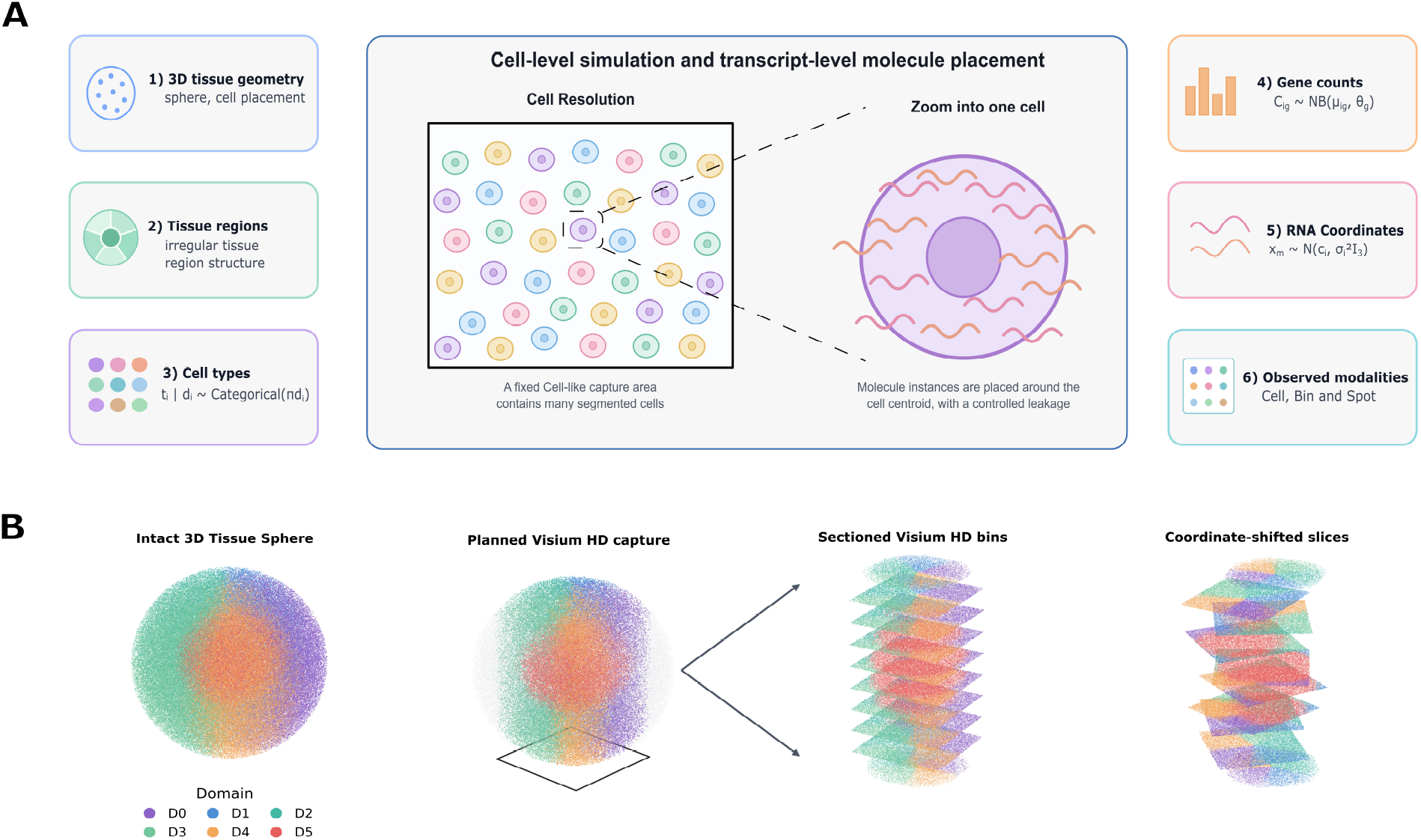
ALBIS is a multi-dimensional, multi-resolution simulator for spatial transcriptomics data. **(A)** Schematic of the six-step generative framework to simulate multi-dimensional and multi-resolution spatial transcriptomics data with ALBIS. Simulation framework includes generating the cell-level 3D tissue coordinates, assigning tissue regions and cell types to simulated cells coordinates, simulating gene expression and transcript-level information, and generating multi-resolution spatial transcriptomics datasets. **(B)** Example of a simulated cell-level sphere that is profiled using 8 × 8 *µm* bins within a 6.5 × 6.5 mm capture window, followed by serial sectioning and independent spatial perturbation of each section to generate misaligned benchmark inputs while retaining known ground-truth coordinates.

### 2.2 ALBIS generates realistic synthetic spatial data across resolutions

Next, we generated synthetic data from ALBIS and compared key statistical properties between the synthetic data to experimentally-derived data across spatial resolutions. Our objective was to demonstrate the flexibility of ALBIS and its ability to be parameterized to approximate experimentally-derived datasets generated using different spatial resolutions, including a cell-, bin-, and spot-level resolution.

We first examined the relationship between gene-level mean expression and variance across experimentally derived and synthetic datasets. Spatial transcriptomics count data are typically overdispersed and are commonly modeled using a negative binomial distribution, making the mean–variance relationship an important statistical property for assessing whether simulated data recapitulate the variability observed in experimental data [18–20]. We additionally compared total counts per observation and the distribution of nonzero expression values before and after preprocessing to assess whether ALBIS reproduced broader characteristics of the count distributions across spatial resolutions. For these analyses, we matched the number of genes across experimental and simulated datasets to ensure comparability of the resulting distributions.

Across spatial resolutions, ALBIS-generated data broadly reproduced the statistical properties of experimentally derived datasets, although the degree of agreement varied by measurement resolution and the statistical property evaluated **(Figure 2; Figure S2**). At cell resolution, simulations parameterized against a single Xenium lung tissue slice recapitulated the characteristic overdispersed mean-variance relationship of the experimental counts, with fitted negative binomial dispersion parameters of 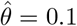 for the simulated cell-resolution data and 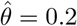 for the corresponding Xenium dataset. Total counts per cell were also closely reproduced (Jensen–Shannon divergence (JSD) = 0.021), although agreement was weaker for the distribution of log-normalized counts (JSD = 0.13).

**Figure 2:**
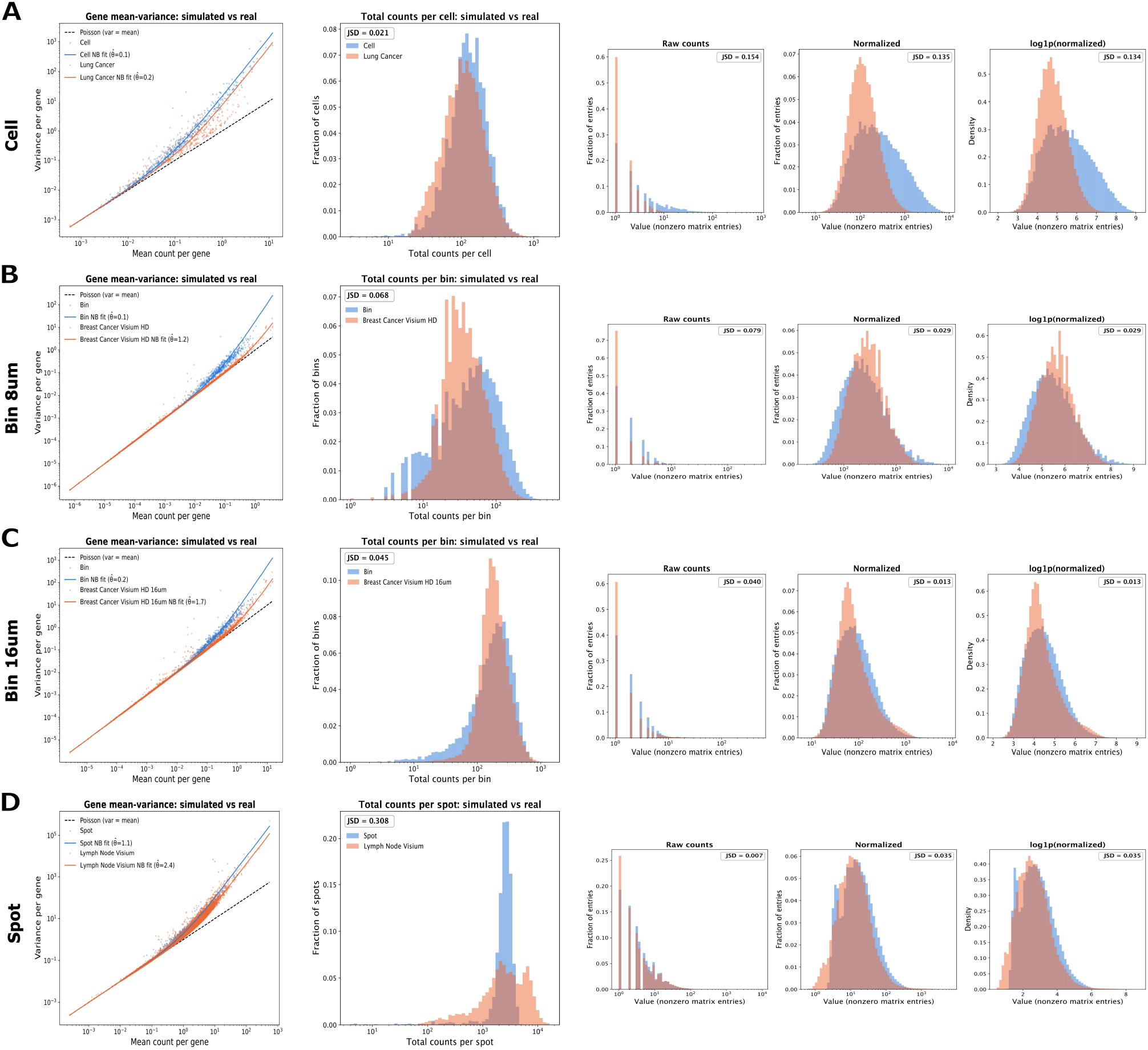
ALBIS generates realistic synthetic spatial data across spatial resolutions. **(A)** Cell-resolution simulations compared with a Xenium lung cancer dataset. **(B)** 8 µm bin-resolution simulations compared with a Visium HD breast cancer dataset. **(C)** 16 µm bin-resolution simulations compared with the corresponding Visium HD breast cancer dataset. **(D)** Spot-resolution simulations compared with a Visium lymph node dataset. For each measurement resolution, the left panel compares the gene-level mean–variance relationship between slice 5 of the simulated data and the real dataset after QC, with fitted negative-binomial relationships and the Poisson expectation (variance= mean). The middle panel compares the distribution of raw total molecular counts per observation. The right panels compare nonzero expression values before normalization, after library-size normalization to 10,000 counts per observation, and after log(1 + *x)* transformation. For the distributional comparisons, the larger gene panel is reduced to its most highly variable genes to match the size of the smaller one. Jensen–Shannon divergence (JSD) is reported for each distributional comparison; lower values indicate greater similarity.

Similar behavior was observed at coarser spatial resolutions. The 8 µm and 16 µm datasets were derived from the same underlying simulation by aggregating observations into grids of different bin sizes. Across both resolutions, the simulated data closely reproduced the statistical properties of the experimental Visium HD breast cancer dataset **(Figure 2B, C)** and, to a lesser extent, those of the pancreatic dataset **(Figure S2B, C)**. Similarly, the spot-level simulations showed close agreement with their experimental counterparts **(Figure 2D Figure S2D)** across gene-level mean-variance relationships, library-size distributions, and expression-distribution summaries. Furthermore these statistical properties are also maintained throughout all 10 simulated slices **(Figure S3)**. Thus, ALBIS can be parameterized to reproduce broad statistical characteristics across cell-, bin-, and spot-level datasets.

### 2.3 Cell type and spatial domain recovery across simulated resolutions

As a next step, we assessed whether ALBIS preserves cell type and spatial domain structure that can be recovered using standard spatial transcriptomics workflows. Cell type recovery was evaluated using PCA, Harmony integration [21], and Leiden [22] clustering, while spatial domain recovery additionally incorporated spatial information using BANKSY [23]. Because cell type and domain labels are retained during simulation, recovered clusters could be directly compared with ground truth using adjusted Rand index (ARI). Clustering analyses in Figure 3 were performed jointly across all 10 simulated tissue sections.

**Figure 3:**
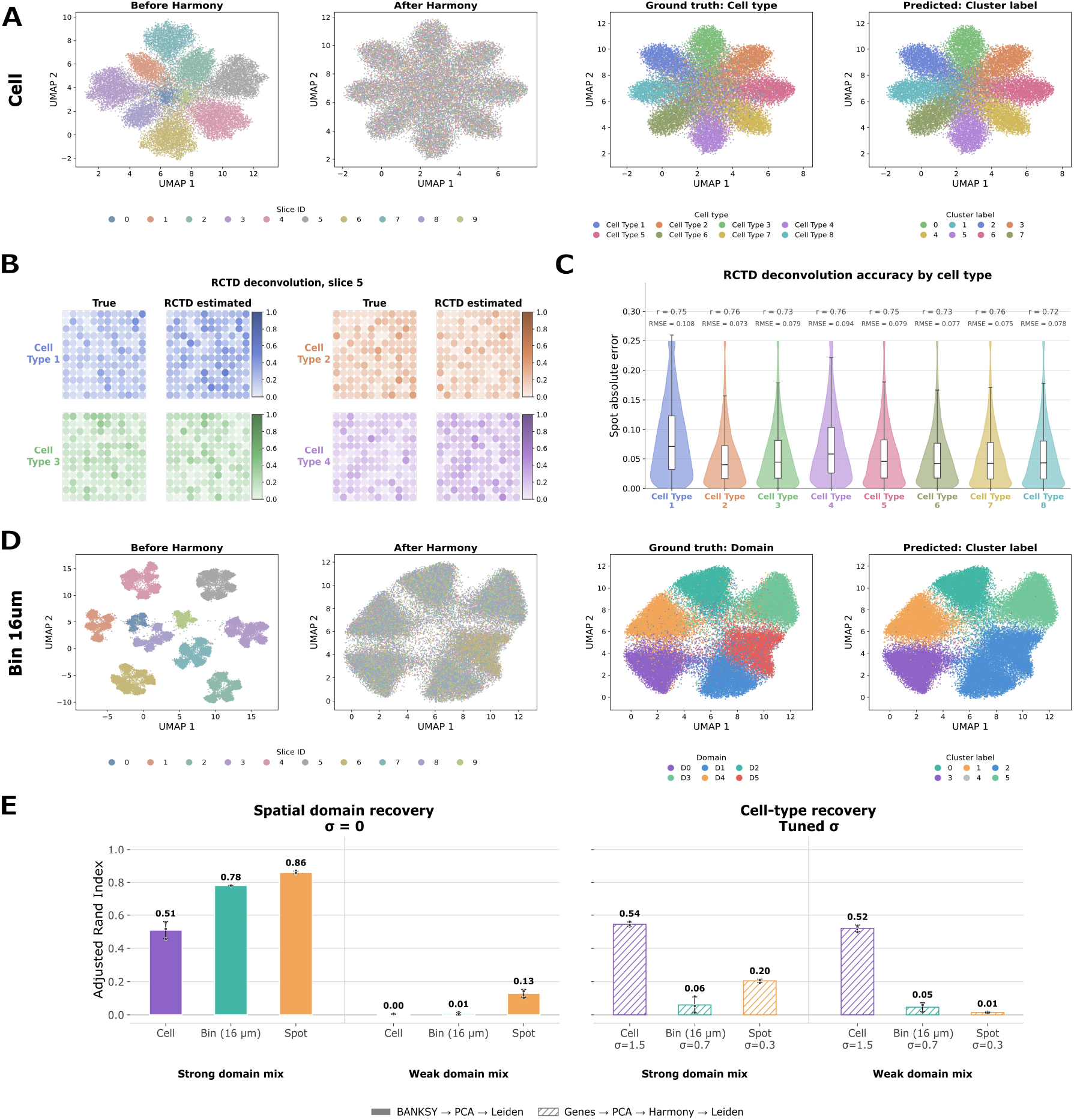
Cell type and spatial domain recovery across simulated resolutions. (A) Cell type clustering at single-cell resolution across 10 simulated tissue sections. UMAP embeddings before and after Harmony integration are colored by simulated slice ID, followed by comparison of simulated ground-truth cell types with Leiden cluster labels. **(B)** Representative RCTD deconvolution results for simulated tissue section 5. True and RCTD-estimated spatial mixture fractions are shown for four representative cell types. RCTD estimates were averaged across three independently generated reference datasets. **(C)** RCTD accuracy across eight simulated cell types, using the average of estimates from three independently seeded single-cell references. Violin plots show the per-spot absolute error of the averaged cell type fraction estimate, with embedded boxplots indicating the median and interquartile range. Pearson *r* and RMSE between averaged estimated and true fractions are shown above each distribution. **(D)** Spatial domain clustering for 16-µm bin-resolution data. UMAP embeddings before and after Harmony integration are colored by simulated tissue section, followed by comparison of known spatial domain labels with Leiden clusters recovered from the BANKSY-derived representation. **(E)** Quantitative recovery of spatial domains and cell types across cell, 16-µm bin, and spot resolutions for strong- and weak-domain-mix simulations. Bars show adjusted Rand index (ARI) between recovered clusters and ground-truth labels using spatially informed BANKSY or expression-only clustering under the indicated batch-effect settings.

At single-cell resolution **(Figure 3A)**, we introduced slice-specific batch effects representing technical shifts in gene expression between tissue sections that were independent of the underlying cell identities. These effects initially drove separation by tissue section in the low-dimensional embedding. Harmony integration substantially reduced this section-associated variation while retaining cell type structure, and Leiden clustering of the integrated representation recovered the simulated cell type labels with an ARI of 0.52 **(Figure 3E)**. Cell type recovery is inherently more challenging at bin and spot resolutions because each observation aggregates transcripts from multiple cell types **(Figure 3E)**. Therefore, we evaluated whether the known cellular mixtures underlying simulated spots could be recovered by spatial deconvolution. Because ALBIS retains the cell type identity of each simulated molecule, the exact ground-truth cell type transcript fractions are known for each spot. RCTD-estimated fractions closely reproduced the spatial distributions of the corresponding ground-truth fractions for representative cell types **(Figure 3B)** and showed generally low per-spot absolute errors across all eight simulated cell types **(Figure 3C)**. Pearson correlations between estimated and true fractions ranged from *r* = 0.72 to 0.76, with RMSE values of 0.073–0.108. Together, these results demonstrate that biological cell type structure remains recoverable from ALBIS-generated data and that its retained ground truth enables direct quantitative evaluation of spatial deconvolution methods.

For spatial domain recovery, the initial simulation setting, which imposed only modest domain-specific differences in cell type composition, did not generate sufficiently distinct spatial structure for reliable domain recovery. We therefore increased the domain-specific differences in cell type composition, which produced more clearly separable spatial domains across single-cell, 16 µm bin **(Figure 3D,E)**, and spot resolutions. Importantly, the strong-domain-mixing setting retained statistical properties comparable to the original simulations **(Figure S4)** and also supported accurate spatial deconvolution at spot resolution **(Figure S5 A, B)** In the absence of an imposed batch effect, domain-recovery ARI reached 0.51, 0.78, and 0.86 for cells, 16 µm bins, and spots, respectively **(Figure 3E)**. As the magnitude of the simulated section-specific batch effect increased, domain recovery generally declined across spatial resolutions **(Figure S5C**). Complete results for clustering, across both datasets and domain vs cell type clustering are available in **(Figure S5D**).

Together, these results show that ALBIS preserves biologically meaningful cell type and spatial domain structure across multiple spatial resolutions while retaining exact ground truth for quantitative evaluation. This enables direct assessment of how well downstream methods recover cell identities, cell type mixtures, and spatial domains, as well as how recovery changes with spatial aggregation and technical batch effects.

### 2.4 Benchmarking spatial alignment and spatial domain recovery with ALBIS

We next demonstrate how the known spatial ground truth generated by ALBIS can be used to benchmark spatial transcriptomics methods across several common analysis settings. Because simulated tissue sections retain known 3D coordinates, spatial domain identities, and matched representations across measurement resolutions, ALBIS enables direct evaluation of spatial alignment, spatial domain recovery, robustness to technical variation, and cross-resolution (from different technologies) integration.

We first evaluated spatial alignment by asking whether STAIR [24] could recover the known geometry of independently perturbed tissue sections **(Figure 4A)**. Across cell-, 16 *µm* bin-, and spot-resolution simulations, STAIR substantially reduced spatial reconstruction error relative to the unaligned input. Because the original coordinates are retained by ALBIS, this improvement could be quantified directly against the true spatial coordinates rather than through an indirect proxy, illustrating how the simulator provides a controlled benchmark for reconstruction accuracy across resolutions.

**Figure 4:**
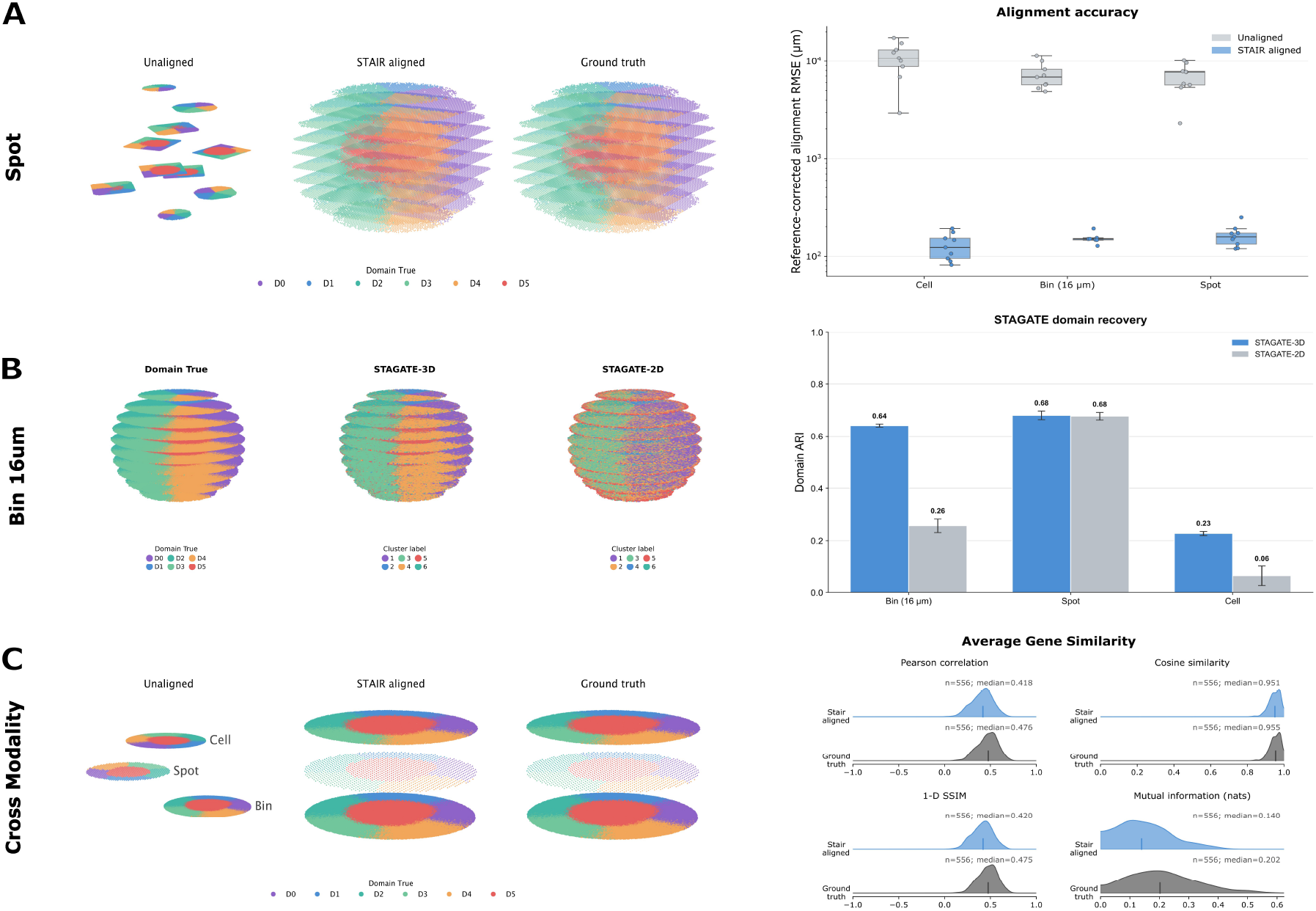
Benchmarking spatial alignment and spatial domain recovery with ALBIS. **(A)** Evaluation of STAIR alignment across cell-, 16 *µm* bin-, and spot-resolution simulations. Spot-sections are shown before alignment, after STAIR alignment, and in the known ground-truth configuration. Accuracy is measured by in-plane RMSE after rigid frame correction using section 0; points represent the nine remaining sections. **(B)** Spatial domain recovery using STAGATE with 2D or 3D spatial graphs. The qualitative example compares inferred clusters with ground truth for the 16 *µm* bin resolution simulation (σ= 0). Recovery is quantified by ARI at bin, spot, and cell resolution. Points represent three independent simulation seeds; lines and error bars show the mean and sample standard deviation. **(C)** Cross-technology alignment of 16 *µm* bin-, spot-, and cell-resolution measurements from the same molecular simulation. Coordinates are shown before alignment, after STAIR alignment, and at ground truth. Gene-level cross-modality similarity is evaluated separately using aligned and ground-truth coordinates. Distributions show Pearson correlation, cosine similarity, one-dimensional SSIM, and mutual information across 556 genes, with each gene’s score averaged over bin–spot and bin–cell comparisons.

We next used ALBIS to benchmark spatial domain recovery with STAGATE [25] under controlled changes in graph construction and technical variation **(Figure 4B)**. Domain recovery was compared using either 2D spatial graphs restricted to within-section relationships or 3D graphs that additionally incorporated cross-section connectivity. Across cell-, 16 *µm* bin-, and spot-resolution simulations, incorporating 3D spatial information produced the largest improvement in domain recovery at the 16 *µm* bin resolution, whereas gains at the spot and cell resolutions were comparatively modest. We further evaluated robustness to increasing slice-specific batch effects. Consistent with the clustering results in **Figure S5E**, domain recovery generally declined as batch-effect magnitude increased, although the extent of this deterioration varied across spatial resolutions and graph-construction strategies. Together, these analyses demonstrate how ALBIS can be used not only to benchmark methods under a single simulation setting, but also to systematically identify the conditions under which their performance begins to degrade.

Finally, we evaluated cross-technology spatial alignment using cell-, 16 *µm* bin-, and spot-resolution measurements generated from the same underlying simulated tissue **(Figure 4C**)-Despite differences in capture resolution, all three modalities share a common molecular ground truth, enabling direct evaluation of cross-resolution alignment. Following independent perturbation and STAIR alignment, the recovered coordinates recapitulated the shared spatial domain organization, while cross-modality gene-similarity distributions approached those observed under the ground-truth configuration. Together, these analyses illustrate how ALBIS can support benchmarking of spatial reconstruction and domain recovery across measurement resolutions, technical perturbations, and cross-technology settings.

### 2.5 Comparison of ALBIS to existing methods

A review of existing spatial transcriptomics simulators **(Table S1)** identified scCube and Spider as methods that, similar to ALBIS, support the generation of 3D spatial transcriptomics data. scCube uses a variational autoencoder (VAE) to generate gene expression profiles and supports user-defined 3D spatial patterns, whereas Spider generates spatial cell arrangements based on user-defined cell type proportions and cell type adjacency relationships. To provide a comparable evaluation with ALBIS, both methods were run reference-free.

We first compared the ability of the three simulators to generate matched 3D tissues across multiple spatial resolutions **(Figure 5A**). For each method, 600,000 cells were generated in three dimensions and divided into 10 tissue sections. The resulting single-cell data were aggregated to generate bin- and spot-resolution measurements. Spider supports geometric specification of the aggregation regions, allowing bin- and spot-level measurements. In contrast, in scCube the aggregation was defined by the number of cells assigned to each bin or spot rather than by a technology-specific capture geometry. By defining the target amount of cells in each spot, scCube will aggregate the cells into spots, without respecting a fixed grid resolution **(Figure S6A)**. Consequently, the resulting spot-level sections did not reproduce the regular spatial arrangement characteristic of Visium arrays.

**Figure 5:**
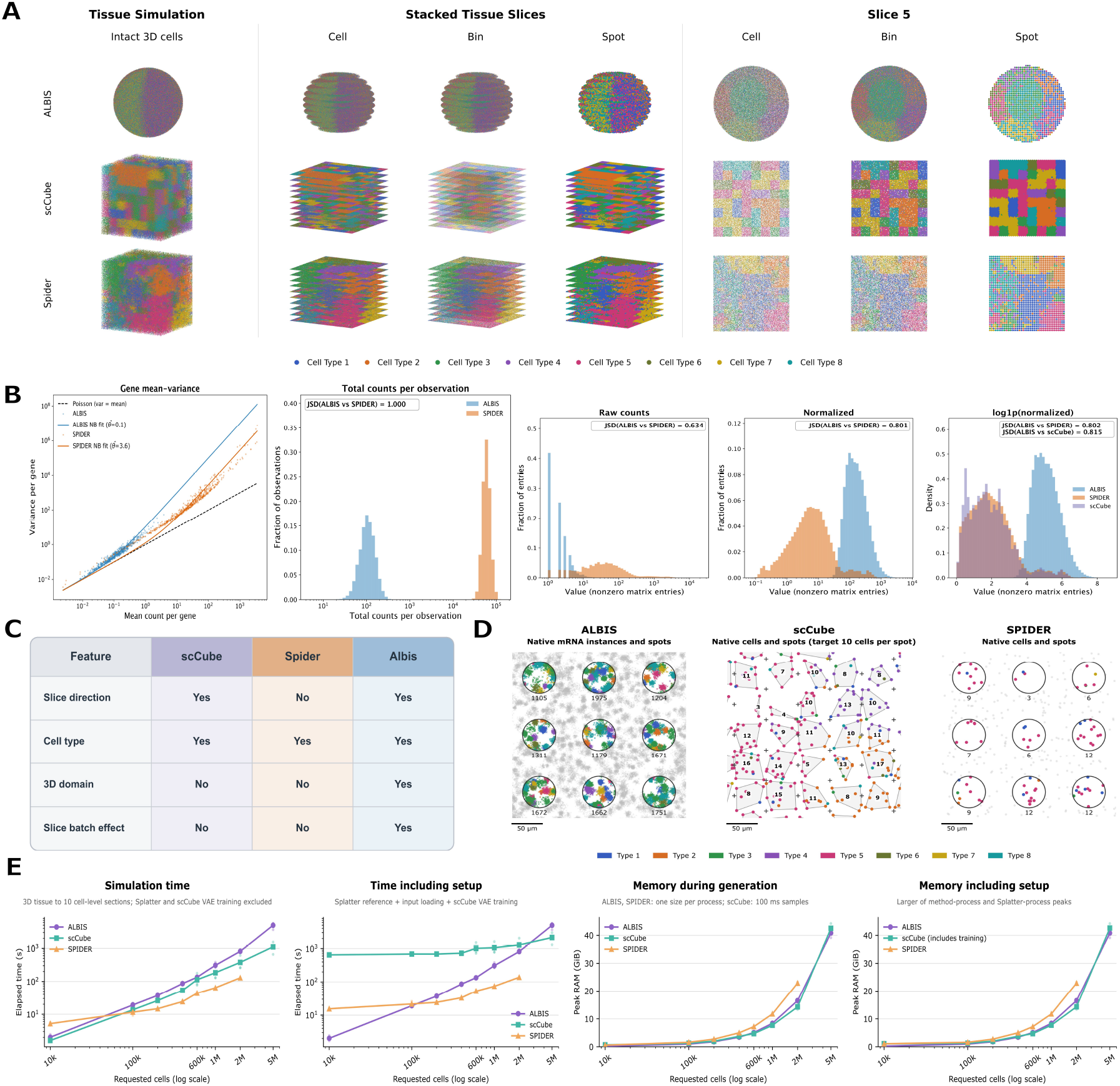
Comparison of ALBIS to existing methods for multi-resolution and multi-dimensional synthetic spatial transcriptomics data. **(A)** Representative spatial layouts generated with ALBIS, scCube, and Spider shown for the full tissue simulation, followed by the stacked tissue slices and a representative slice (slice 5) at cell-, bin-, and spot-level resolution. **(B)** Comparison of cell-level expression properties, including total counts per cell, gene mean–variance relationships, and raw, normalized, and log-normalized expression distributions. **(C)** Comparison of supported simulation features, including slice direction, cell type, 3D tissue regions, and slice-specific batch effects. **(D)** Comparison of the underlying spatial representation. ALBIS simulates individual mRNA molecules (crosses), with 55 µm spots capturing only transcripts within their boundaries, including those from cells that lie partly inside the spot and from transcript spillover. scCube and SPIDER generate cell-level profiles (dots): SPIDER assigns cells to circular spots, whereas scCube assigns each cell to a square grid tile. scCube’s reported spot centres (+) are offset by half a spacing in alternate columns to mimic Visium and may fall outside their assigned cells. Numbers indicate transcripts per spot (ALBIS) or cells per spot (scCube, SPIDER). **(E)** Computational performance across increasing simulation sizes, comparing generation and setup-inclusive runtime and peak memory usage. SPIDER did not complete at 5,000,000 cells: sectioning its expression exceeded the 32-bit index limit of scipy’s sparse matrices, independent of available memory.

We next compared the statistical properties of simulated expression data across cell, bin, and spot resolutions. Because the scCube reference-free workflow used here generates expression values on a transformed rather than raw-count scale, comparisons involving scCube were restricted to log-normalized expression values. At the cell level, ALBIS and Spider showed similar gene mean–variance relationships, while their raw expression distributions differed, with ALBIS exhibiting a greater proportion of low-count observations **(Figure 5B**). This difference remained apparent after normalization and log transformation, with the ALBIS distribution shifted toward higher expression values. At the bin and spot resolutions, however, the distributions generated by ALBIS and Spider became more similar, particularly after normalization and log transformation **(Figure S6B**,**C)**.

ALBIS also differs from scCube and Spider in the spatial features represented by the simulation. In the configurations evaluated here, scCube and Spider do not natively provide 3D spatial domain annotations or explicitly model slice-specific batch effects, while both are supported by ALBIS **(Figure 5C)**. A further distinction is the underlying simulation unit **(Figure 5D**). Whereas scCube and Spider generate expression profiles at the cell level, ALBIS explicitly simulates the spatial coordinates of individual mRNA molecules. This molecule-level representation enables properties such as transcript spillover to be modeled directly, allowing molecules originating from one cell to be captured within a neighboring cell or spatial observation.

Finally, we compared computational requirements across methods over increasing simulation sizes **(Figure 5E**). Generation times were broadly comparable, although their scaling differed with increasing cell number. Memory consumption was similar across methods, with Spider requiring slightly more memory at larger sizes and failing to complete the five-million-cell run.

## 3 Discussion

In this work, we demonstrate that ALBIS can generate spatial transcriptomics data at multiple resolutions from the same underlying set of simulated molecular transcripts, preserving coordinates across cell-, bin-, and spot-level representations. The framework is highly customizable, allowing biological and technical parameters to be tuned to reproduce key statistical properties of experimental datasets. We further show that simulated biological structure is recoverable using commonly used spatial transcriptomics workflows, with cell types recovered across resolutions and spatial deconvolution accurately estimating cell type proportions at the spot level.

Spatial transcriptomics is increasingly moving beyond individual 2D tissue sections toward the integration and reconstruction of spatial information across tissue sections and measurement resolutions [26−28]. However, experimental datasets often lack known spatial coordinates across sections and resolutions, making simulation an important tool for controlled benchmarking. In this context, ALBIS differs from existing 3D spatial simulators in its molecule-level representation of the underlying tissue. scCube [17] and Spider [5], for example, support both reference-based and reference-free simulation but generate expression at the cell level before aggregation into bins or spots. ALBIS instead simulates spatially positioned mRNA molecules from which cell-, bin-, and spot-level measurements are derived, providing a common ground truth across resolutions. This is particularly relevant for benchmarking cross-resolution alignment and integration methods, where coordinates between different representations of the same tissue is itself part of the task being evaluated. Furthermore, by simulating individual molecules, ALBIS can explicitly model transcript spillover, whereby a configurable fraction of transcripts is displaced from its originating cell into the surrounding tissue. Such effects represent an intrinsic source of spatial mixing in experimentally generated spatial transcriptomics data [29] and can therefore be directly controlled and evaluated within the ALBIS framework.

However, ALBIS also has several limitations in its current implementation. First, ALBIS currently simulates only transcriptomic measurements and does not model additional molecular modalities such as protein abundance or chromatin accessibility. Extending the framework to multimodal spatial assays could enable benchmarking of methods designed for joint analysis across molecular layers. Second, ALBIS does not currently generate paired histological or fluorescence images, such as H&E or immunofluorescence images, which limits its use for evaluating methods that integrate spatial transcriptomics with tissue morphology or image-derived features. Third, spatial domains in ALBIS are currently defined primarily through differences in cell type composition. The framework does not yet model domain-specific transcriptional programs within the same cell type or spatially variable genes that arise independently of compositional differences. In experimental tissues, such variation can reflect differences in cellular state and the local microenvironment [30]. Incorporating these effects could provide a more realistic setting for evaluating methods that identify tissue regions from both compositional and transcriptional differences. Finally, ALBIS currently operates exclusively in a reference-free setting, whereas several existing spatial transcriptomics simulators can use experimental references to inform simulated gene expression and spatial organization. Extending ALBIS with a reference-based mode could improve its ability to reproduce the characteristics of specific experimental tissues.

Overall, we envision ALBIS as a flexible framework for developing and benchmarking computational methods for spatial transcriptomics, particularly those that rely on shared information across resolutions or tissue sections. ALBIS could also support agentic benchmarking [31], providing controlled datasets for evaluating how AI agents select, configure, and execute analysis workflows and assess their results. Its ability to generate spatially related sections with known molecular, cellular, spatial-domain, and 3D ground truth supports the evaluation of methods for alignment, integration, clustering, deconvolution, and reconstruction.

## 4 Methods

### 4.1 Overview of the simulator

The simulator generates a synthetic 3D tissue volume, assigns cells to spatial domains and cell types, simulates gene expression counts, expands counts into individual RNA molecules, and then derives multiple spatial transcriptomics measurements from the same underlying molecular ground truth. This design allows the same simulated tissue to be observed at cellular resolution, subcellular bin-level resolution, or multicellular spot-level resolution. Because all simulated measurements are generated from the same 3D transcript-level ground truth, the simulator provides known ground-truth information for evaluating coordinate recovery, domain preservation, cell type mixture recovery, and cross-resolution correspondence.

#### 3D tissue geometry and cell placement

We modeled the tissue as a 3D spherical volume

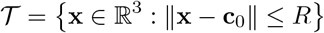

where **c**_**0**_ is the tissue center and *R* is the tissue radius. Cell centroids were sampled uniformly within this sphere. Each cell *i* was assigned a centroid c_*i*_ ∈ ℝ^3^, a radius *r*_*i*_, a spatial domain label *d*_*i*_, and a cell type label *t*_*i*_. Cell radii were sampled from a truncated log-normal distribution.

#### Tissue Regions with irregular boundaries

To create a spatial structure that was more challenging than simple layered or regular geometric domains, we defined multiple 3D tissue domains with irregular boundaries. The default simulated tissue contained four domains. One domain represented a central core, while the remaining domains formed angular wedge-like regions around the core. Rather than using perfectly smooth boundaries, the simulator introduced boundary irregularity through smooth sinusoidal noise fields. The core boundary was perturbed by direction-dependent 3D smooth noise, producing a bumpy core surface, and wedge boundaries were perturbed by angular noise, producing wavy domain interfaces. These perturbations create realistic imperfect domain boundaries while preserving a known 3D ground truth.

#### Cell type composition by domain

Cell types were sampled conditional on spatial domain. For each domain *d*, we defined a categorical distribution over *K* cell types,

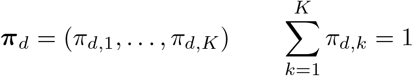

Given a domain label *d*_*i*_, the cell type of cell *i* was sampled as

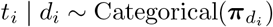

The default parameters simulate 4 cell types. Domain-specific cell type mixtures were designed so that the wedge domains were each enriched for one dominant cell type (up to 70% of that domain’s composition, with the remaining three types making up the rest), while the central core domain contained a uniform mixture of all cell types. This design avoids unrealistically pure domains and creates a benchmark in which alignment methods must preserve both spatial structure and local cell type composition that varies systematically between core and peripheral regions.

#### Gene programs and per-cell gene expression

For each cell type, marker genes were defined together with shared marker genes and noise genes. In the default parameters, each cell type was assigned 80 marker genes, with 25% of marker genes shared across cell types and approximately **10%** additional noise genes. Let *m*_*t,g*_ denote the multiplicative effect of cell type *t* on gene *g*. Cell type specific marker genes were assigned larger expression multipliers, shared marker genes were assigned intermediate multipliers, and noise genes were assigned weaker or nonspecific expression effects.

For each gene *g*, a baseline expression level *λg* was sampled from a log-normal distribution. Each cell *i* was assigned a cell-specific size factor *S*_*i*_, and optional domain-level scale factors were allowed. The expected expression of gene *g* in cell *i* was modeled as

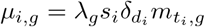

where 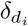 is a domain-specific scale factor and 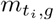 is the cell type specific gene multiplier. Counts were then sampled using the NB2 negative binomial parameterization,

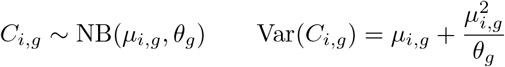

where *θg* is the gene-specific inverse-dispersion parameter. In the main simulation, the inverse-dispersion parameter was centered at *θ* = 25, with gene-specific jitter.

#### Transcript-level molecule simulation

After simulating the cell-by-gene count matrix, we expanded each count 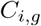 into individual RNA molecule instances. For each cell *i*, the total number of molecules was

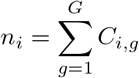

Each molecule inherited the source cell identity, source cell type, source domain, and gene label. This transcript-level representation provides the common ground truth from which all downstream measurement resolutions were generated.

For each molecule generated from cell *i*, a 3D coordinate was sampled from an isotropic Gaussian distribution centered at the source cell centroid,

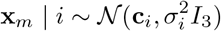

The standard deviation *σ*_*i*_ was chosen so that 95% of molecules were expected to fall within the spherical cell boundary of radius *r*_*i*_. Since

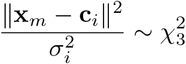

we set

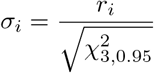

where 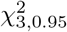 is the 95th percentile of the chi-square distribution with three degrees of freedom. This construction allows a controlled leakage tail, with approximately 5% of molecules falling outside the source cell boundary. This leakage is useful for mimicking segmentation uncertainty, transcript diffusion, or local molecule assignment ambiguity in imaging-based spatial transcriptomics.

#### Cell-level observation and spillover

To mimic technical limitations in cell segmentation and transcript assigment, cellular-resolution observations were intentionally generated based on observed molecule coordinates rather than source cell identity. A molecule at location *X*_*m*_ was assigned to the cell whose spherical boundary contained it. Molecules that did not fall inside any cell boundary were treated as unassigned for cellular-resolution observations. The observed count for cell *j* and gene *g* was therefore

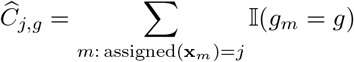

Because assignment depends on observed molecule location, a molecule generated from source cell *i* can be counted in a different cell *j* if its coordinate falls inside cell j’s boundary. This creates controlled transcript spillover while preserving the source-cell ground truth.

#### Multi-resolution measurements from the same transcript-level truth

The same simulated molecule set was used to derive multiple spatial transcriptomics resolutions. For the cell-level representation, molecule counts were aggregated to segmented cells as described above. For the bin-level representation, intended to approximate VisiumHD-like data, the 3D tissue volume was sliced along a specified axis and molecules were aggregated into regular square bins on the corresponding two-dimensional slice plane. For the spot-level representation, intended to approximate Visium-like data, molecules were assigned to regularly spaced circular capture spots within each slice plane. This multi-resolution design is the main distinction between our simulation framework and perturbation-based benchmarks that only transform a single observed dataset.

For bin-level data, each molecule was assigned to a slice index *s* and a two-dimensional bin *p*. The bin-level count matrix was defined as

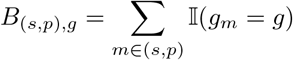

For spot-level data, regularly spaced capture spots were placed in each slice plane, and molecules were assigned to the nearest spot if their in-plane distance was within the capture radius. The spot-level count matrix was defined as

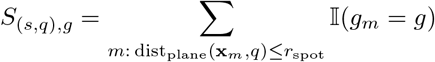

Because each molecule retains its source cell type and source domain, the ground-truth cell type and domain mixture for every bin or spot can be computed directly. For example, the true cell type fraction for bin *(s,p)* and cell type *k* is

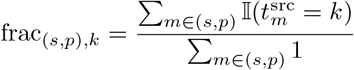

#### Platform-specific capture windows

To model differences in field of view across spatial transcriptomics technologies, different capture windows were applied to the same simulated tissue. In the default configuration, the cell-level Xenium-like representation used a conceptual capture region of 12 × 24 mm, while the Visium-like and Visium HD-like representations used a 6.5 × 6.5 mm capture window. The Visium HD-like bin size was set to 8 *µm*, and the Visium-like spot grid used 100 *µm* spacing with a 27.5 *µm* capture radius. These values were chosen to approximate representative settings of the corresponding technologies, but the capture area, bin size, spot spacing, and capture radius are user-configurable and can be modified according to the intended simulation setting. These settings allow the simulator to generate observations with different spatial resolutions and capture areas from the same underlying 3D tissue.

#### Slicing, batch effects, and unaligned coordinates

In order to create multiple slices, by default, the simulated 3D tissue is cut into 5 equal-width sections along each of *x, y*, and *z* axes. This produced separate sectioned datasets for each slicing direction and allows for the evaluation of algorithms under different anatomical orientations. To mimic slice-to-slice technical variation, we apply a gene-wise multiplicative batch effect for each slice:

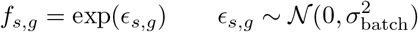

Given an observed count 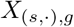 the batch-affected count was generated by Poisson resampling:

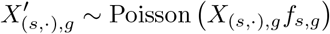

This procedure preserved nonnegative count values while introducing slice-specific gene shifts. To create misaligned inputs for alignment benchmarking, each slice was also perturbed by an independent in-plane rigid transformation. By default in main simulation, rotations were sampled up to 180° and translations up to 200 *µm*. The simulator stored both the aligned ground-truth coordinates and the perturbed unaligned coordinates, enabling direct comparison between method outputs and known ground truth.

### 4.2 Statistical Properties of the Simulated Data

We next evaluated whether the simulated datasets reproduced key statistical properties observed in real spatial transcriptomics data at the corresponding Xenium-like, Visium HD-like, and Visium-like measurement resolutions. We described quality control of the real datasets in **Section 4.6** For the simulated datasets, we removed observations with `total_counts = 0` or fewer than three detected genes prior to analysis and utilized slice 5 of the sphere.

We focused on three properties of the count distributions. First, we compared the gene-level mean–variance relationship, with particular emphasis on the dispersion parameter 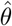, to assess whether the simulated data reproduced the overdispersion observed in the corresponding real datasets. Second, we compared the distributions of total counts per observation. Third, we compared the distributions of nonzero expression values using raw counts, after librarysize normalization and after a subsequent log(1 + *x)* transformation. For gene-matched comparisons, we reduced the dataset with the larger gene panel to the size of the smaller one by selecting its top highly variable genes with Scanpy’s `highly_variable_genes` (`flavor=“ seurat_ v3”`, applied to raw counts of the QC-filtered data); this sub-setted the real Visium and Visium HD panels to 556 genes, and the simulated panel to the 392 genes of the Xenium reference.

#### 4.2.1 Empirical Dispersion Estimation

For each simulated and real dataset, we estimated a dataset-level dispersion parameter, 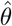, from the corresponding quality-controlled, raw counts, post-batch count matrix. We calculated the estimate over the full gene panel, independently of the HVG-matched comparisons. For each gene *g*, we calculated the empirical mean, 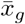, and population variance, *v*_*g*_, across observations.

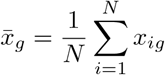

and the population variance,

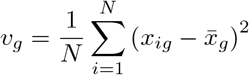

across observations. Under the NB2 parameterization,

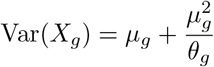

the gene-specific method-of-moments estimate was

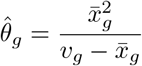

We calculated this estimate only for genes with 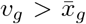, since genes with variance less than or equal to the mean did not yield a finite positive dispersion estimate under this parameterization. We summarized the dataset-level dispersion as the median across eligible genes,

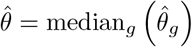

We then plotted the corresponding negative-binomial mean-variance relationship,

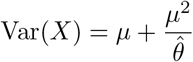

alongside the empirical gene-level means and variances. We included the Poisson relationship, Var(*X*) = *µ*, as a reference.

#### 4.2.2 Distributional Similarity

We quantify agreement between simulated and real distributions using Jensen–Shannon divergence (JSD), computed from normalized histograms with 60 shared bin edges spanning the pooled range. JSD was computed as the square of the Jensen–Shannon distance returned by `scipy. spatial. distance. Jensenshannon` (SciPy vl.15.3) with base-2 logarithms, bounding JSD between 0 (identical distributions) and **1**.

We evaluate total counts and detected genes per observation, and nonzero expression values in raw, library-size-normalized, and log(1 + x)-transformed matrices. Nonzero values are randomly subsampled to at most 2 × 10^6^ per dataset using a fixed seed before binning.

#### 4.2.3 Simulator tuning

We iteratively tune the generative parameters for each modality by generating simulated data and comparing them with the corresponding real dataset. We primarily tune the simulator to match the gene-level mean-variance relationship and the estimated dispersion 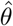. We additionally compare the distributions of total counts, detected genes, and nonzero expression values across raw, library-size-normalized, and log(1 + *x*)-transformed data, using JSD as a quantitative measure of distributional similarity.

Based on these comparisons, we adjust the parameters controlling dispersion *θ*, dispersion jitter, baseline expression and cell count. For the aggregated modalities, we also adjust tissue-packing density. We repeat this process until the simulated data reasonably reproduce the corresponding real-data distributions across the different preprocessing stages. The parameters contributing most strongly to the fit differ across modalities. Cell-resolution data are primarily influenced by the underlying count distribution, whereas bin- and spot-resolution data are additionally affected by spatial aggregation and tissue packing. We therefore tune each modality independently. The final parameter configurations are reported in **Supplementary Table S2**.

### 4.3 Clustering Pipeline

We evaluated recovery of the simulated cell types and spatial domains. All analyses processed the ten slices jointly, after removing observations with zero total counts or fewer than three detected genes. For conditions with batch effects, we integrated sections with Harmony (v2.0.0) [21], treating each section as a batch.

#### Leiden clustering

In every workflow, we built a 15-nearest-neighbor graph using 30 principal components (Harmony-corrected where applicable) across all sections and applied Leiden clustering (python-igraph v0.11.9, via Scanpy v1.11.5). To compare recovery at the known number of ground-truth categories, we tuned the resolution by bisection over [0.01, 3] for at most 15 steps, targeting eight cell types or six spatial domains. Runs that did not reach the target cluster count were scored at the resolution yielding the closest number of clusters; they were excluded from the bar-panel means but retained in the batch-strength sweep. For bins and spots, the ground-truth cell-type label was the dominant cell type of each observation.

We quantified recovery using the adjusted Rand index (ARI), comparing the inferred clusters with the simulated labels pooled across all ten sections:

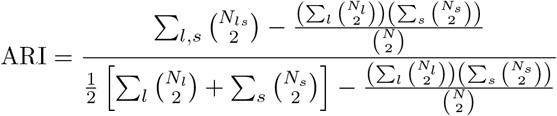

Here, *N* is the total number of observations, 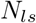 is the number assigned to inferred cluster land ground-truth category *s*, and *N*_*1*_ = ∑ *L*_*s*_ *N*_*1s*_ and *N*_*s*_ = ∑_l_ *N*_*1s*_ are the corresponding totals. An ARI of 1 indicates perfect agreement, values near 0 indicate agreement expected by chance, and negative values indicate agreement below chance.

#### 4.3.1 Cell type recovery

We used the simulated datasets of Figure [2], generated with the baseline domain-by-cell-type composition matrix, in which domains differed only modestly in cell type composition, and repeated the analysis on the domain-enriched simulation described below, each for simulation seeds 2025, 101, and 202. We fixed batch-effect strengths per modality: σ_batch_ = 1.5 for cells, 0.7 for 16-µm bins, and 0.3 for spots. We normalized counts to 10^4^ per observation, log(1 + x)-transformed them, and scaled them per gene; we ran PCA on all genes and corrected the PCs with Harmony. We followed this with Leiden clustering.

#### 4.3.2 Spatial domain recovery

For spatial domain recovery, we used a dedicated domain-enriched simulation with stronger coupling between spatial domain membership and cell type composition. We assigned each of the six domains two signature cell types with sampling probabilities of approximately 30% each, while each of the remaining six cell types had a probability of approximately 6.7%. The domain-enrichment setting changed the domain by cell type sampling probabilities while retaining the same spatial-placement and gene-expression generation procedures. We built neighborhood-augmented expression matrices with BANKSY (v0.0.7) [23] from counts normalized to 10^4^ per observation (no log transform), offsetting slice coordinates so that neighborhoods did not cross sections. The parameter *λ* weighted an observation’s own expression *(λ* = 0) against its neighborhood *(λ* = 1), and *k*_geom_ set the number of spatial neighbors: 60, 100, and 8 for cells, 16-µm bins, and spots, which corresponded to different physical distances across modalities.

##### Batch-zero comparison

At σ_batch_ = 0 we ran BANKSY → PCA → Leiden without Harmony. In addition to spatial domains, we used the same pipeline to recover cell types, testing whether neighborhood information helped or hurt this non-spatial target. Clustering targeted six clusters for domains and eight for cell types. We selected *λ* on a single seed (2025) of the domain-enriched simulation, sweeping λ ∈{O, 0.1, 0.2, 0.3, 0.5, 0.8, 1.0} and choosing, separately for each target, the *λ* with the highest ARI among runs reaching the target cluster count (ties favored smaller. λ): 0.3, 0.3 and 0.5 for domains and 0.1, 0.1 and 0 for cell types (cells, bins, spots). We applied these values unchanged to the baseline (weak) domain mixture and ran the same batch-zero comparison on both mixtures.

##### Batch-strength sweep

On the domain-enriched simulation we ran BANKSY → PCA →Harmony → Leiden across increasing σ_batch_, with kgeom as above and the domain *λ* selected in the batch-zero comparison (0.3, 0.3 and 0.5) and reported the mean and standard deviation across simulation seeds. All three modalities used the same σ_batch_ grid: 0 to 0.5 in steps of 0.05, plus 0.6, 0.7, 1.0 and 1.5 (15 values). We held parameters fixed across all batch strengths, and applied Harmony at every point, including σ_batch_ = 0. We retained all runs and reported the ARI at the resolution whose cluster count was closest to the target, including runs where the target was not reached.

#### 4.3.3 Spatial deconvolution with RCTD

We evaluated spot-level cell type deconvolution using RCTD (Robust Cell Type Decomposition; [32])through `spacexr v2.2`. **1** in `full` mode, allowing multiple contributing cell types per spot. The query comprised 10,033 simulated Visium-like spots across all ten slices, with 100 µm spacing and a 27.5 µm capture radius. The single-cell reference contained 24,207 cells, eight ground-truth cell types, and the same 556-gene panel. We generated both datasets with the Figure 2 simulation parameters and the baseline domain-cell type composition setting **(Supplementary Table S2)**, including their modality-specific batch effects (σ_batch_ = 1.5for the reference and 0.3 for the query). RCTD received raw counts and total UMI counts, with cell type labels we supplied for the reference; we did not use reference coordinates.

We defined ground-truth spot compositions as the fractions of captured molecules contributed by each cell type. We row-normalized estimated weights to sum to one and compared them with ground truth for the 9,896 spots (98.6%) passing RCTD’s quality-control and minimum-UM! filtering; the same spots passed filtering in every run. To assess sensitivity to reference variation, we ran RCTD with three references: the canonical reference, generated with the same seed as the query, and two references generated from independent seeds, holding all other parameters and the query fixed. We then averaged the three estimated fraction matrices elementwise to obtain a reference-averaged estimate. We quantified performance using Pearson correlation and RMSE pooled across all spot–cell type pairs, per spot across cell types, and per cell type across spots.

### 4.4 Applications of ALBIS

#### 4.4.1 Alignment Error with STAIR

We first evaluated ALBIS as a benchmark for spatial reconstruction, using STAIR (v1.3.1) [24] to align the 10 sections of a simulated z-stack. STAIR combines gene-expression information across sections and performed rigid alignment through mutual-nearest-neighbor initialization followed by iterative closest-point refinement. ALBIS independently perturbed each section by an in-plane rotation and translation, and we withheld these parameters from STAIR. We captured the same simulated tissue (sphere radius 4,600 µm, 10^6^ cells) at three resolutions: Visium HD-like 16 µm bins, Visium-like spots and single cells.

Because each reconstruction had an arbitrary global frame, section 0 served as a fixed reference. For each condition (unaligned or STAIR-aligned), we fitted a rigid transformation (rotation and translation; no scaling or reflection) from the ground-truth coordinates of section 0 to its coordinates in that condition. We then applied this transformation to all other sections, with no section-specific fitting. For each non-reference section *s*, alignment error was

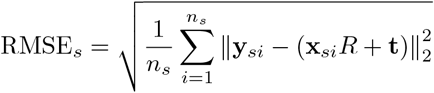

where *X*_*si*_ and **y**_*si*_ were the ground-truth and evaluated (unaligned or aligned) coordinates of observation *i*, and *R*, **t** were the condition-specific reference transformation. We knew observation correspondence from the simulation, so we did not need nearest-neighbor matching. Excluding the reference section gave nine RMSE values (in µm) per condition.

#### 4.4.2 Spatial Clustering with STAGATE

Second, we evaluated ALBIS for benchmarking 3D spatial domain identification using STAGATE (v1.0.0) [25] a graph attention autoencoder that learns a low-dimensional representation from expression and a spatial neighbor graph. We compared two formulations, both trained on the pooled ten-section z-stack using the aligned simulated coordinates: a 2D formulation with within-section edges only, and a 3D formulation that additionally connected observations in adjacent sections. For bins and spots, edges connected observations within 1.5 times the median nearest-neighbor spacing (24 µm and 150 µm, respectively), both within and between sections; single-cell graphs used six nearest neighbors within each section and three nearest neighbors in each adjacent section, queried in both directions. This comparison tested whether cross-section information improved recovery of the known spatial domains.

We applied both formulations to Visium HD–like 16 µm bins, Visium-like spots and single cells, using the strong domain-composition setting in which the six ground-truth domains had distinct cell-type enrichments. We varied per-section batch effects across *σ* = 0, *σ* = 0.05 and the modality-specific value used in the batch-correction analysis *(σ* = 0.7 for bins, 0.3 for spots and 1.5 for cells), without explicit batch correction. We used all 556 genes after library-size normalization and log transformation, and trained STAGATE for 500 epochs with the training seed fixed at 0. We evaluated each condition on three simulation seeds.

We clustered each embedding with `mclust` into six components, matching the number of ground-truth domains. We quantified domain recovery by the adjusted Rand index (ARI) between inferred and ground-truth domain labels, computed once per replicate and condition over all retained observations pooled across the ten sections. We reported results as mean ARI ± sample standard deviation across simulation seeds, with individual replicate values overlaid.

#### 4.4.3 Cross-Modality Alignment with STAIR

As a third application, we evaluated cross-modality alignment. We captured a single simulated tissue, generated with the 16 µm bin expression parameters, as 16 µm bins, Visium-like spots and single cells; we independently rigidly perturbed each modality’s copy of section 5 and then aligned it with STAIR.

We evaluated cross-modality expression similarity with the bins as the reference. We normalized counts to 10,000 per observation and log-transformed them, and computed the mean expression of each of the 556 shared genes in square grids (200 µm for bin–spot, 300 µm for bin–cell). We retained grids if both modalities had at least half the lower of their two mean per-grid occupancies. We computed Pearson correlation,

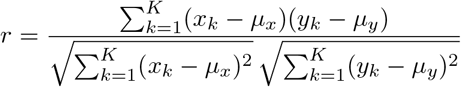

cosine similarity,

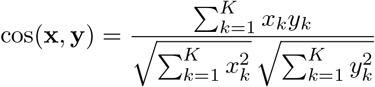

one-dimensional structural similarity (SSIM),

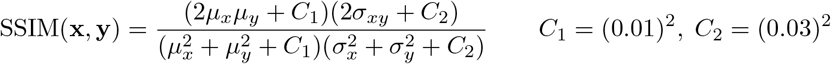

and mutual information,

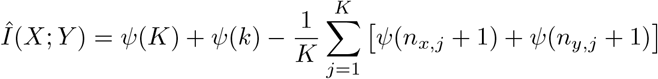

across retained grids, including zero-expression values, and averaged them per gene over the two modality pairs, requiring valid scores for both. We compared distributions between STAIR-aligned and ground-truth coordinates, after a rigid frame correction estimated from the bins.

### 4.5 Comparison to other Spatial ‘Iranscriptomic Simulation Methods

We compared ALBIS with spatial transcriptomics simulators that could generate a 3D tissue and section it: scCube [17] and Spider [5]. Each method simulated 600,000 cells and 556 genes within the same 4.1 mm extent (a cube for scCube and Spider; a sphere of radius 2,050 µm for ALBIS), divided into ten sections and captured at cell, bin and spot resolution using each method’s own functions. Because both methods required external expression input in their reference-free modes, we generated a shared synthetic expression pool with Splatter (v1.26.0) [6]: 10,000 cells, 556 genes and eight equally sized groups corresponding to cell types.

scCube (v2.0.0) [17] supports reference-based and reference-free simulation; we used the reference-free mode. A variational autoencoder (VAE) generates gene-expression profiles; we could train it on reference data or load it from a pretrained model. To avoid observed reference data, we trained the VAE (200 epochs) on the Splatter pool, which scCube library-size normalized and log-transformed internally, and generated 600,000 cells in equal cell-type proportions. scCube imposed spatial organization separately. Of scCube’s reference-free spatial generators, only the random-pattern generator supported three dimensions. The random-pattern generator placed cells uniformly at random, sampled a Gaussian random field on a coarse grid (8 × 8 × 8; *δ* = 2) and assigned cell types to grid units by thresholding the field at quantiles matching the cell-type proportions; with λ= 0. 75, it then shuffled the labels of a random 25% of cells. We scaled its unitless coordinates to the 4.1 mm cube. We divided the volume into ten sections with scCube’s splitting option, and applied scCube’s two-dimensional aggregation to each section to obtain bin- and spot-level outputs. Rather than using a physical capture size, scCube set the capture-grid spacing from a target mean number of cells per unit (three per bin and ten per spot here, giving units of approximately 29 µm and 53 µm on square and hexagonal layouts, respectively) and summed the expression values of the cells assigned to each unit.

Spider (st-spider v1.2.0) [5] also supports reference-based and reference-free simulation; we used the reference-free setting, in which an external simulator, here the Splatter pool, supplied expression profiles. Spider places 600,000 cells uniformly within the volume and assigned cell types by matching a user-specified cell-type transition matrix with simulated annealing, first on a coarse spatial grid and then at progressively finer scales down to individual cells. We specified equal cell-type proportions and a same-type neighbor probability of 0.7, with the remaining probability distributed equally among the other cell types. We then assigned each cell an expression profile sampled with replacement from the Splatter cells of its type. We divided the volume into ten equal-thickness sections with Spider’s slicing function, and generated bins (16 µm squares) and spots (55 µm diameter circles on a square 100 µm grid) per section with Spider’s native aggregation, which summed the counts of the cells in each capture unit.

For the comparison of statistical properties, we followed the methodology described in Section 4.2, applied to the fifth section of each method’s output. Because we trained scCube’s VAE on log-normalized expression values and it generated these values rather than raw counts, we excluded scCube from count-based statistics. At bin and spot resolution, scCube’s values were sums of cell-level log-expression values and we therefore showed them separately.

We benchmarked ALBIS, scCube and SPIDER at eight sizes from 10,000 to 5,000,000 cells (556 genes, one CPU thread, three simulation seeds per method and size) on shared cluster nodes of a single hardware type (Intel Xeon Platinum 8558U). Each method ran through its native API, and each size ran as a separate program. Tissue extent (and, for ALBIS, sphere radius) was scaled from the 600,000-cell configuration by (n/600,000)^1^/^3^, keeping cell density constant. Simulation time was the wall-clock time from building the 3D tissue to cell-level data in ten sections along *z*. Setup-inclusive time added Splatter reference generation and input loading (scCube, SPIDER) and VAE training (scCube; 200 epochs on the 10,000-cell reference, retrained in every run, no checkpoints saved); ALBIS required no setup.

Memory was each program’s peak RAM use. For scCube, generation memory excludes VAE training and was sampled every 100 ms. Setup-inclusive memory is the most RAM needed at any point in the full workflow: the larger of the Splatter reference’s and the method’s peak, as the two run one after the other. We report the mean over seeds and show each seed individually.

### 4.6 Overview of Data

To assess how realistic our simulations were, we compared the cell-level, bin-level, and spot-level outputs with real datasets from each technology. Here we briefly described the quality control (QC) we applied to these real datasets; we showed QC results in **Supplementary Figure S1**. We removed flagged observations, and applied no normalization before comparison.

#### 4.6.1 Healthy and diseased lung, Xenium

For the Xenium samples, we flagged low-quality cells using a median absolute deviation (MAD)-based approach: we flagged cells more than 4 MADs below the median of log1p-transformed total transcript counts or number of detected genes. We additionally flagged cells with zero count density (transcripts per unit cell area) and cells with any control-probe or control-codeword counts.

#### 4.6.2 Breast cancer and pancreas (Visium HD); lymph node and tonsil (Visium)

For Visium HD and Visium data, we performed QC with SpotSweeper’s [33,34] local outlier detection on in-tissue observations, at 8 µm and 16 µm bins for Visium HD and at spot resolution for Visium. For each observation we calculated total UMI counts, number of detected genes and the percentage of counts from mitochondrial genes, and identified local outliers for each metric relative to its 36 nearest neighbors using a robust z-score cutoff of 3. We log-transformed total counts and detected genes and flagged them in the lower tail; we flagged mitochondrial percentages in the upper tail.

### 4.7 Data availability

The raw spatial transcriptomics datasets used in this study are publicly available from 10x Genomics:

1. [Human Breast Cancer](https://www.10xgenomics.com/datasets/visium-hd-cytassist-11mm-human-breast-cancer), generated using Visium HD.
2. [Human Pancreas](https://www.10xgenomics.com/datasets/visium-hd-cytassist-11mm-human-pancreas), generated using Visium HD.
3. [Human Lung and Diseased Lung](https://www.10xgenomics.com/datasets/xenium-human-lung-preview-data-1-standard), generated using Xenium.
4. [Human Lymph Node](https://www.10xgenomics.com/datasets/visium-v2-human-lymph-node), generated using Visium.
5. [Human Tonsil](https://www.10xgenomics.com/datasets/visium-cytassist-gene-and-protein-expression-library-of-human-tonsil-with-add-on-antibodies-h-e-6-5-mm-ffpe-2-standard), generated using Visium.

## 4.8 Code availability

Code to reproduce all preprocessing, analyses, and figures in this manuscript is available on GitHub at https://github.com/JanMatthias1/albis_paper. A python implementation is available on PyPI at https://pypi.org/project/ALBIS/0.1.1/ and on our GitHub at https://github.com/JanMatthias1/sim_app/

## 4.9 Acknowledgments, Funding, Authorship Contributions

## Acknowledgments

We acknowledge the Joint High Performance Computing Exchange (JHPCE) at the Johns Hopkins Bloomberg School of Public Health for computational support, and members of the Hicks lab for their valuable feedback. Furthermore we acknowledge the use of Artificial Intelligence to refine the language of the manuscript and assist with coding. The authors take full responsibility for its content and integrity.

## Author Contributions Statement

- Conceptualization: JM, JY, SCH
- Methodology: JY, JM
- Investigation and review of existing papers: JM
- Data aggregation and preprocessing: JM
- Formal analysis: JM
- Writing – original draft: JM, JY, SCH
- Writing – review & editing: JM, JY, SCH
- Visualization: JM
- Supervision: SCH
- Funding acquisition: SCH
- Project administration: SCH

