## Supplementary Figures and Tables for "ALBIS: simulation of multi-resolution and multi-dimensional spatial transcriptomics data"

### Contents

1. **Supplemental Tables**
   (a) Supplemental Table S1: Comparison of spatial transcriptomics simulation frameworks
   (b) Supplemental Table S2: Generative simulation parameters
2. **Supplemental Figures**
   (a) Supplemental Figure S1: Quality control filtering
   (b) Supplemental Figure S2: Library-size distributions
   (c) Supplemental Figure S3: Statistical properties across spatial resolutions
   (d) Supplemental Figure S4: Weak- versus strong-domain-mix simulations
   (e) Supplemental Figure S5: Cell type and spatial domain recovery

**Supplementary Table S1:** Comparison of spatial transcriptomics simulation frameworks. Tissue dimensionality distinguishes 2D spatial simulation, 3D cell-layout generation, and reference-guided interpolation between sections.

| Method | Primary representation | Reference mode | New layout | Tissue dimensionality | Modeled effects and distinguishing features |
|---|---|---|---|---|---|
| Dissociated scRNA-seq simulators[a] | Cell-by-gene count matrices | Varies; no ST reference | No | Not applicable; no spatial tissue | Count-level biological and technical variation without a native spatial model. |
| simAdaptor[b] | Cluster-wise counts placed at reference coordinates | ST reference-based | No | One reference 2D section at a time | Reference locations are clustered, each cluster is simulated independently, and the resulting counts are returned to the original spatial coordinates. |
| *Related or task-specific spatial simulation workflows* | | | | | |
| stLearn [35] | Cell- or spot-level expression and neighbourhood scenarios | scRNA-seq or ST based | Yes | One 2D spatial layout at a time | Analysis-specific simulations of cellular neighbourhoods and cell–cell-interaction scenarios. |
| spaSim [36] | Spatial cell-point patterns and tissue-like images | Reference-free | Yes | One 2D spatial field at a time | Models cell density, clustering, colocalization, and tissue-like spatial patterns, but does not generate a complete gene-expression count matrix. |
| scSpatialSIM [37] | Single-cell point patterns with categorical or continuous molecular attributes | Reference-free | Yes | One 2D spatial field at a time | Models tissue compartments, holes, cell clustering, colocalization, and expression gradients; primarily intended for spatial point-pattern benchmarking. |
| scMultiSim [38] | Single-cell expression, multiomic profiles, and spatial cell layouts | Reference-free | Yes | One 2D spatial map at a time | Provides gene-regulatory-network and cell–cell-interaction ground truth together with biological and technical variation. |
| SpatialcoGCN-Sim [39] | Spatially assigned scRNA-seq cells aggregated into spots | ST and scRNA-seq based | Yes[c] | One reference-derived 2D section at a time | Uses a VAE and nearest-neighbour mapping to infer cell coordinates; supports adjustable spot resolution, cell mixtures, and neighbour-derived expression noise. |
| spatialSimGP [40] | Poisson gene counts at supplied spatial locations | Reference-free | User supplied | One 2D spatial layout at a time | Uses a gene-specific Gaussian process to generate spatial expression; supports spatial and non-spatial genes, configurable spatial variance, and fixed or gene-specific correlation length scales. |

[a] Splatter [6], ZINB-WaVE [7], scDesign2 [8], SPARSim [9], and SymSim [10]. [b] simAdaptor [11] clusters an ST reference with BayesSpace [12] and simulates each cluster independently while retaining the reference coordinates. [c] SpatialcoGCN-Sim generates new cell and spot coordinates, but their spatial organization is learned from a reference ST section.

**Supplementary Table S1:** Comparison of spatial transcriptomics simulation frameworks (*continued*).

| Method | Primary representation | Reference mode | New layout | Tissue dimensionality | Modeled effects and distinguishing features |
|---|---|---|---|---|---|
| *General-purpose spatial gene-expression simulators* | | | | | |
| SRTsim [13] | Gene counts at spatial locations | Both | Yes | One 2D tissue section at a time | Poisson, NB, ZIP, or ZINB counts with tissue- or domain-level spatial patterns; values at new locations can be informed by neighbouring reference locations. |
| scDesign3 [14] | Counts conditioned on spatial coordinates and covariates | Reference-based | Limited[d] | One 2D tissue section at a time | Flexible marginal models and copulas; supports spatial, batch, condition, cell type, and other continuous or discrete covariates. |
| sCCIgen [41] | Cell profiles, optionally aggregated into multicell regions | Spatial reference optional [g] | Yes | One 2D spatial map at a time | Models cell colocalization, regional expression effects, and spatially dependent cell–cell and gene–gene interactions. |
| SimSpace [42] | Spatial cell layouts, cell types, and modular omics profiles | Both | Yes | 2D spatial maps or 3D cell layouts[e] | Uses a Markov random field to model tissue organization, spatial niches, cell type colocalization, density, and local interactions. |
| FEAST [16] | Gene-expression profiles in simulated or interpolated spatial slices | Reference-guided | Yes | 2D slice simulation with reference-guided 3D interpolation and reconstruction[f] | Models gene-level mean, variance, and sparsity; supports controlled transcriptional changes, spatial-pattern perturbations, virtual-slice generation, and geometric transformations for alignment benchmarking. |
| scCube [17] | Simulated cell- or spot-level expression profiles | Both[g] | Yes | 2D or 3D cell patterns; generated 3D patterns can be divided into serial 2D slices | VAE-based expression generation with reference-based or reference-free spatial patterns, adjustable slice number, spot arrangement, resolution, and gene panels. |
| Spider [5] | Single-cell profiles, optionally aggregated into spots | Spatial reference optional [g] | Yes | 2D or 3D cell layouts; generated 3D layouts can be divided into sequential 2D slices | Models cell type proportions and neighbourhood transition patterns; supports customizable tissue domains, spot geometry, and multi-slice organization. |
| **ALBIS** | **Cells → individual 3D RNA molecules → cells, bins, and spots** | **Reference-free** | **Yes** | **One continuous 3D tissue, sectioned along the $x$, $y$, or $z$ axis** | **Models cell-type-dependent expression and domain-specific cell type composition, individual RNA coordinates, serial sections, slice-specific batch effects, molecule-level spillover, platform capture geometry, and rigid coordinate perturbations.** |

[d] Lower-level scDesign3 functions accept new covariate matrices, but the method does not provide a primary de novo tissue-geometry generator. [e] SimSpace simulates 3D cell layouts with cell-type-dependent expression, but it provides no native function to export serial sections as separate datasets. [f] FEAST generates virtual sections by reference-guided interpolation between spatial sections, rather than by sectioning a natively simulated 3D tissue volume. [g] A spatial reference is optional, although an expression reference, trained model, or externally generated expression profiles may still be required.

**Supplementary Table S2: Generative simulation parameters used for comparison with experimental spatial transcriptomics data in Figure 2.** Parameters are shown for cell-, 8 $\mu$m bin-, 16 $\mu$m bin-, and spot-resolution simulations.

<table>
<tr><th>Parameter</th><th>Description</th><th>Cell</th><th>Bin (8 $\mu$m)</th><th>Bin (16 $\mu$m)</th><th>Spot</th></tr>
<tr><td>Aggregation unit</td><td>Spatial unit represented by each observation</td><td>Single cell</td><td>8 $\mu$m bin</td><td>16 $\mu$m bin</td><td>100 $\mu$m spot spacing<br>27.5 $\mu$m capture radius</td></tr>
<tr><td>$n_{\text{cells}}$</td><td>Simulated cells before aggregation</td><td>24,207</td><td></td><td>600,000</td><td></td></tr>
<tr><td>Sphere radius $R$</td><td>3D tissue radius ($\mu$m)</td><td colspan="4">2050</td></tr>
<tr><td>3D packing fraction</td><td>Fraction of sphere volume occupied by cells</td><td>$\sim$0.17%</td><td>$\sim$4%</td><td>$\sim$4%</td><td>$\sim$4%</td></tr>
<tr><td>Capture window</td><td>Instrument capture area</td><td>$12 \times 24$ mm</td><td>$6.5 \times 6.5$ mm</td><td>$6.5 \times 6.5$ mm</td><td>$6.5 \times 6.5$ mm</td></tr>
<tr><td>Platform</td><td>Simulated measurement platform</td><td>Xenium</td><td>Visium HD</td><td>Visium HD</td><td>Visium</td></tr>
<tr><td>Cell radius (mean)</td><td>Mean cell radius ($\mu$m)</td><td colspan="4">7.5</td></tr>
<tr><td colspan="6">Expression model</td></tr>
<tr><td>$\log \mu$</td><td>Lognormal scale parameter for per-gene baseline expression</td><td>$-2.5$</td><td>$-2.5$</td><td>$-2.5$</td><td>$-2.25$</td></tr>
<tr><td>$\log \sigma$</td><td>Scale of per-gene baseline expression</td><td>0.7</td><td>0.7</td><td>0.7</td><td>1.0</td></tr>
<tr><td>NB dispersion $\theta$</td><td>Negative-binomial dispersion (lower = noisier)</td><td>0.40</td><td>2.0</td><td>2.0</td><td>0.25</td></tr>
<tr><td>$\theta$ jitter</td><td>Per-gene variation in $\theta$</td><td>0.15</td><td>0.6</td><td>0.6</td><td>0.10</td></tr>
<tr><td>Marker fold-change</td><td>Expression multiplier for cell type markers</td><td colspan="4">3.5</td></tr>
<tr><td>Shared-marker fold-change</td><td>Expression multiplier for shared markers</td><td colspan="4">2.5</td></tr>
<tr><td>Batch effect $\sigma$</td><td>SD of per-slice, per-gene log-fold-change</td><td>1.5</td><td>0.7</td><td>0.7</td><td>0.3</td></tr>
<tr><td colspan="6">Tissue structure</td></tr>
<tr><td>Tissue slices</td><td>Number of simulated sections</td><td colspan="4">10</td></tr>
<tr><td>Spatial domains</td><td>Number of ground-truth domains</td><td colspan="4">6</td></tr>
<tr><td>Cell types</td><td>Number of ground-truth cell types</td><td colspan="4">8</td></tr>
<tr><td>Gene panel size</td><td>Number of simulated genes</td><td colspan="4">556</td></tr>
</table>

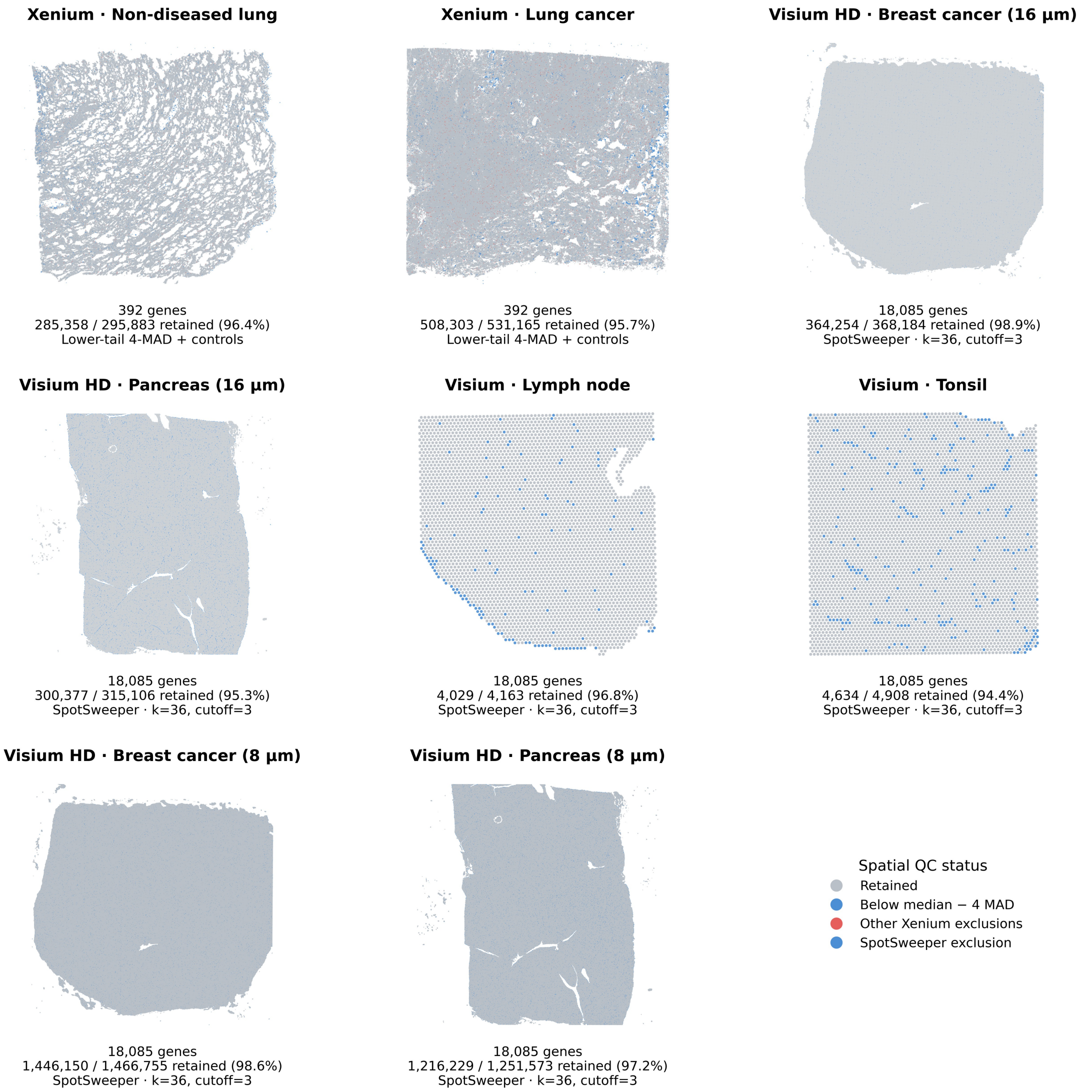


**Supplementary Figure S1: Quality control filtering of experimentally derived spatial transcriptomics datasets used for Figure 2.** 2D spatial plots of retained and filtered observations are shown for (**A**) Xenium non-diseased lung, (**B**) Xenium lung cancer, (**C**) Visium HD breast cancer at 16 $\mu$m resolution, (**D**) Visium HD pancreas at 16 $\mu$m resolution, (**E**) Visium breast cancer, (**F**) Visium lymph node, (**G**) Visium tonsil, (**H**) Visium HD breast cancer at 8 $\mu$m resolution, and (**I**) Visium HD pancreas at 8 $\mu$m resolution. Retained observations are shown in gray, whereas observations removed during quality control filtering are shown in blue and, for the Xenium datasets, additional observations removed by dataset-specific filtering criteria are shown in red. The number and percentage of observations retained after filtering are indicated below each panel. Xenium datasets were filtered using a lower-tail median absolute deviation criterion on total counts, whereas Visium and Visium HD datasets were filtered using SpotSweeper [33, 34] with the thresholds indicated below each panel.

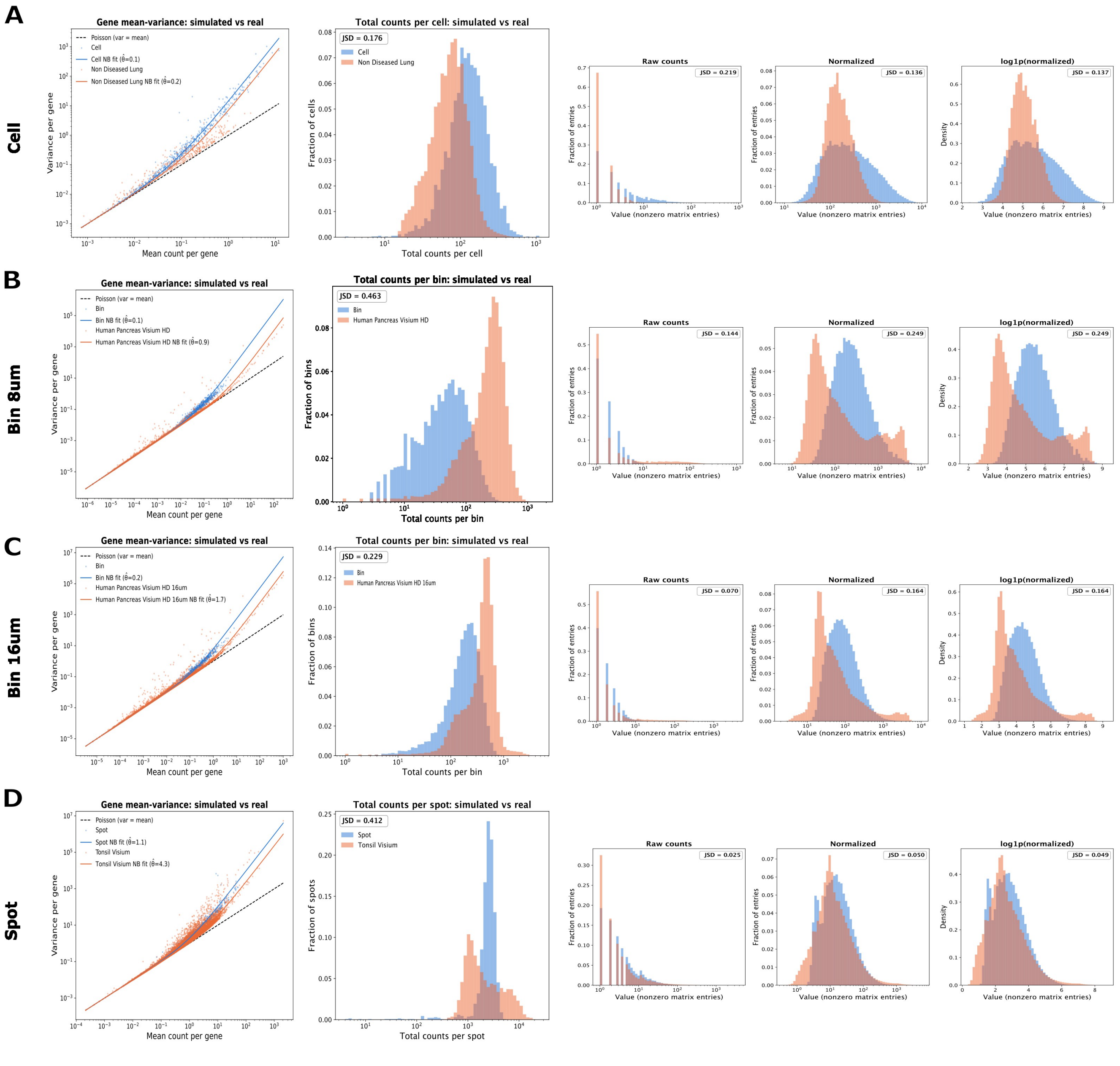


**Supplementary Figure S2: Comparison of library-size distributions between simulated and real-world spatial transcriptomics data.** Comparison of simulated and corresponding real datasets at three measurement resolutions. For each modality, the panels show the gene mean–variance relationship (left), total counts per observation (middle), and nonzero expression values as raw counts, normalized counts and log1p-normalized counts (right). (**A**) Cell-resolution simulated data compared with real-world Xenium data. (**B,C**) Comparison of simulated bin-resolution data with real Visium HD data at 8 and 16 µm bin resolutions. (**D**) Spot-resolution simulated data compared with real-world Visium data.

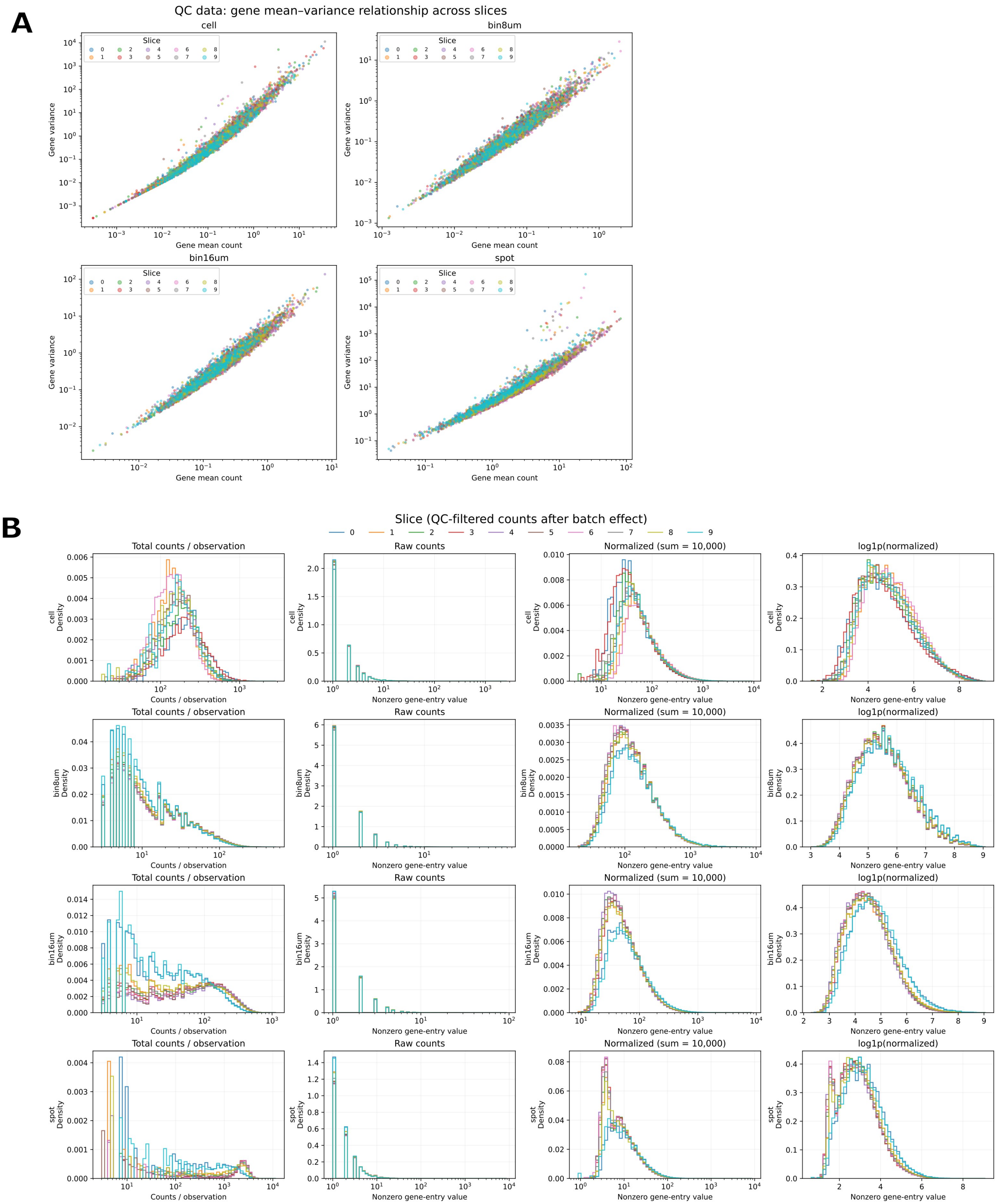


**Supplementary Figure S3: Statistical properties of ALBIS-simulated data across spatial resolutions.** (**A**) Gene-level mean–variance relationships across the 10 simulated tissue sections at cell, 8 µm bin, 16 µm bin, and spot resolution. Each point represents a gene, with colors indicating the simulated tissue section. (**B**) Distributional properties of the simulated data at the same four resolutions. From left to right, panels show total counts per observation, nonzero raw gene-expression values, nonzero expression values after library-size normalization to 10,000 counts per observation, and log1p-transformed normalized expression values. Distributions are shown separately for each simulated tissue section.

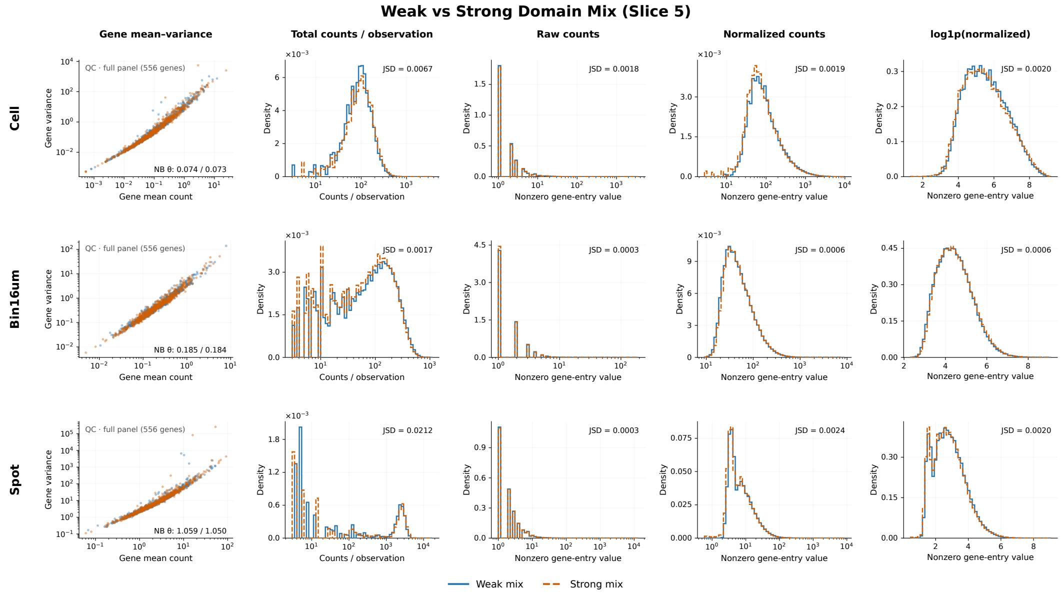


**Supplementary Figure S4: Comparison of statistical properties between weak- and strong-domain-mix ALBIS simulations.** Statistical properties of the weak-domain-mix (blue) and strong-domain-mix (orange) simulations are compared for a representative tissue section (slice 5) at cell, 16 µm bin, and spot resolutions. From left to right, panels show the gene-level mean–variance relationship, total counts per observation, and distributions of nonzero expression values for raw, normalized, and log1p-transformed normalized counts. Negative-binomial dispersion estimates ($\theta$) are reported for the mean–variance comparisons, and Jensen–Shannon divergence (JSD) quantifies differences between the corresponding distributions. Despite the stronger association between spatial domains and cell type composition, the two simulation configurations retain highly similar expression and count-distribution properties across all three spatial resolutions.

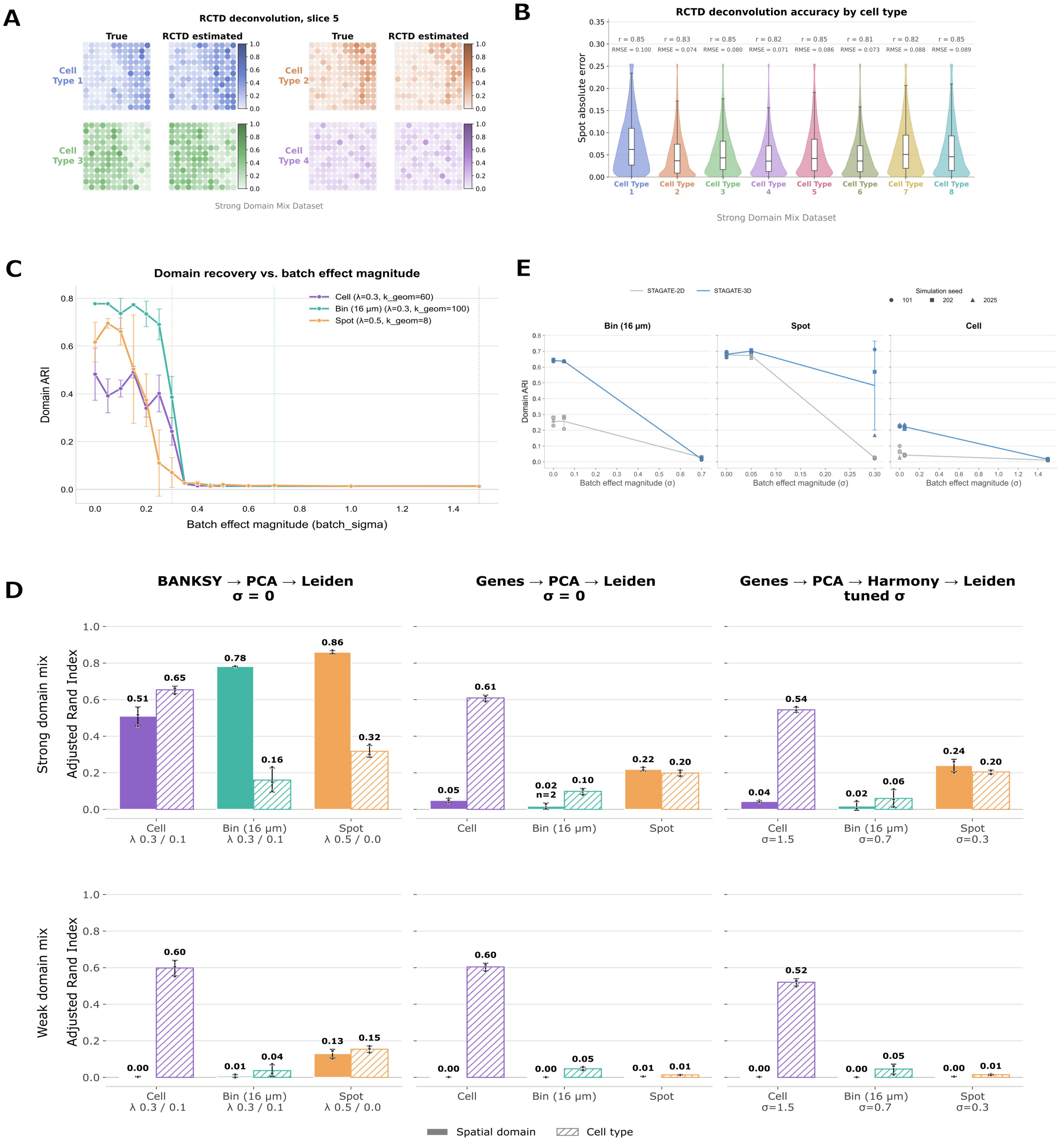


**Supplementary Figure S5: Evaluation of cell type and spatial domain recovery in ALBIS-simulated data.** (**A**) Ground-truth and RCTD-estimated cell type fractions for four representative cell types in slice 5. (**B**) RCTD deconvolution accuracy across all eight cell types, shown as per-spot absolute errors, with Pearson correlation coefficients ($r$) and RMSE reported for each cell type. (**C**) Spatial domain recovery across increasing batch-effect magnitudes at cell, 16 μm bin, and spot resolutions, quantified using adjusted Rand index (ARI). (**D**) Comparison of spatial domain and cell type recovery for strong- and weak-domain-mix simulations using BANKSY-based and expression-only clustering workflows. Results are shown without batch effects and, for the expression-only workflow, at selected batch-effect magnitudes. (**E**) Comparison of STAGATE spatial domain recovery using 2D and 3D spatial graphs across increasing batch-effect magnitudes at 16 μm bin, spot, and cell resolutions. Points indicate independent simulation seeds.

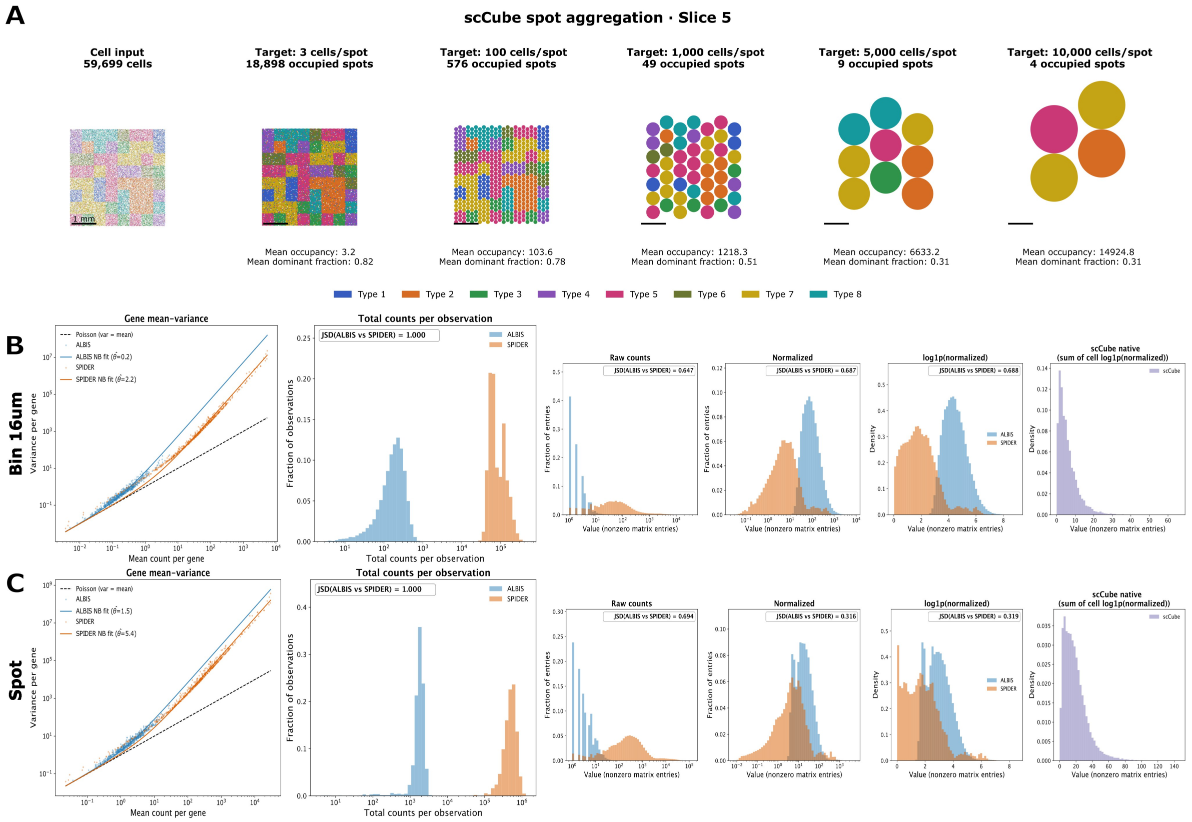


**Supplementary Figure S6: Comparison of simulated spatial transcriptomics data generated by ALBIS, scCube, and SPIDER.** (**A**) Aggregation of scCube-simulated cells into spot-level observations for representative target occupancies ranging from 3 to 10,000 cells per spot. For each aggregation level, the number of occupied spots, mean cell occupancy, and mean dominant cell type fraction are shown. (**B**) Statistical comparison of ALBIS and SPIDER at the 16 µm bin resolution. Shown are gene-level mean–variance relationships with fitted negative-binomial (NB) dispersion estimates, distributions of raw total counts per observation, and distributions of nonzero expression values as raw counts, normalized expression and log1p-transformed normalized expression. Jensen–Shannon divergence (JSD) is reported between ALBIS and SPIDER. Because scCube generates log-normalized cell-level expression rather than raw counts, its aggregated bin-level expression is shown separately. (**C**) Corresponding statistical comparison at the spot resolution.
